# Single-cell mass cytometry maps the evolution of immune cell signatures predictive of acute graft-versus-host disease

**DOI:** 10.1101/2025.02.26.640331

**Authors:** James A Aries, Sarah Charrot, Symeon Theocharidis, Megha Meena, Wing-Yiu Jason Lee, Monica Escorcio Correia, Daniel J Pennington, Jamie Cavenagh, John G. Gribben, Jeff Davies

**Author notes:** Corresponding author: Dr Jeff Davies Address: Centre for Haemato-Oncology, Barts Cancer Institute, Queen Mary University of London, Charterhouse Square, London EC1M 6BQ. **Competing Interest Statement**: The authors have no competing interests to disclose.

## Abstract

Allogeneic haematopoietic stem-cell transplantation (AHST) can cure patients with many diseases, but harmful acute graft-versus-host disease (aGvHD) remains a challenge. Many immune cells are implicated in the pathogenesis and control of alloreactive T-cell responses causing aGvHD, but the functionally dominant cells at different times remain unknown. Using mass cytometry to simultaneously assess alloreactive and immunoregulatory cell populations after AHST, we identified a robust early immune signature predictive of aGvHD rich in CD4 effector memory T-cells and deficient in functionally allosuppressive NK cells. Additionally, network analysis identified a heterogeneous immunoregulatory cell group (IRCG) that more accurately predicted aGVHD than any individual cell populations or established serum biomarkers. Immune signatures preceding aGvHD evolved over time, with qualitative and quantitative changes in both the alloreactive T-cell compartment and the constituent components of the IRCG. In mapping how alloreactive and immunoregulatory cell populations evolve temporally, we provide mechanistic insight into dynamic control of alloreactivity supporting development of time-sensitive targeted strategies to reduce aGvHD.

**Graphical Abstract:** 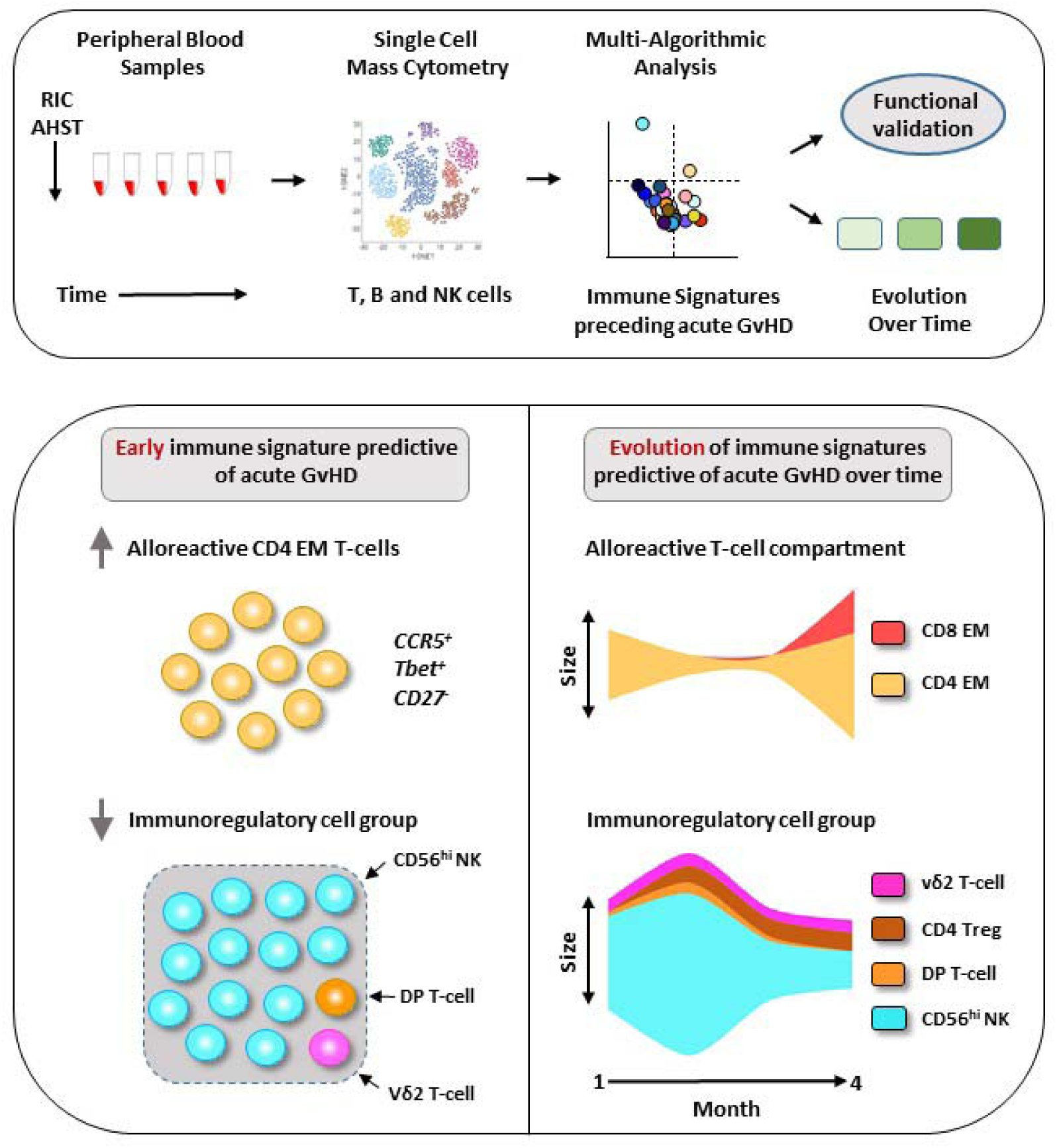

## Introduction

Allogeneic hematopoietic stem-cell transplantation (AHST) is the only curative therapy for many patients with blood cancers and other conditions. Allogeneic donor grafts also contain donor immune cells, some of which can recognize genetically disparate recipient histocompatibility antigens. The resulting alloreactive T-cell response may damage healthy patient tissues resulting in acute graft-versus-host disease (aGvHD), a major barrier to successful AHST.(1–3) Despite the introduction of reduced-intensity conditioning (RIC) which reduces AHST toxicity, aGvHD remains a significant challenge.(4)

Both CD4 and CD8 donor T-cells contribute to the pathogenesis of aGvHD.(5–8) Similarly, several distinct immunoregulatory cell populations have been implicated in the control of alloreactive T-cell responses including CD4 regulatory and CD4/8-double-positive (DP) T-cells, subsets of NK cells, invariant NKT and γδ T-cells.(9–18) However, as most studies focussed on cellular immune profiles at the onset of aGvHD, the temporal dynamics of immune signatures preceding aGvHD have not been clearly defined. Furthermore, integrated studies simultaneously measuring multiple immune cell subsets are lacking but could shed light on the functional dominance of individual alloreactive effector and immunoregulatory cell populations after AHST, leading to more focused approaches to prevent or treat aGvHD.

In recent years mass cytometry (MC) has facilitated simultaneous measurement of multiple immune cell paprameters.(19) Furthermore, new multi-algorithmic and unsupervised approaches to analyse high-dimensional data can better characterize complex immune cell landscapes. Several early studies employing MC have demonstrated the potential of these new approaches to identify cellular immune signatures after AHST but thus far have studied small patient numbers and heterogeneous of transplant platforms and focused on immune cell landscapes at or after the onset of aGvHD.(20–22)

We therefore used single cell MC and multi-algorithmic analytic approaches to identify cellular immune signatures preceding aGvHD in blood of patients from a prospective cohort undergoing RIC-AHST using a uniform transplant platform.(23, 24) We compared the predictive value of early immune signatures to that of established serum biomarkers and mapped the evolution of these signatures and their constituent components over time.

## Materials and Methods

### Ethical approval

The study was approved by the London Research Ethical Committee (05/Q0605/140 and 06/Q0604/110) and conducted in accordance with the Declaration of Helsinki. Patients gave informed consent.

### Patient cohort and AHST platform

We prospectively studied a cohort of consecutive adult patients with hematologic malignancies undergoing RIC-AHST. Patients received uniform conditioning followed by peripheral blood stem-cells from HLA A-, B-, C-, DRB1- and DQ-matched donors without T-cell depletion, and post-transplant immunoprophylaxis with calcineurin inhibition (CNI) and short methotrexate.(24).Peripheral blood samples were collected at up to 5 time-points after AHST up to D+180 or the onset of aGvHD or disease relapse.

### Single-cell MC

A primary panel was constructed consisting of 24 antibody-metal conjugates (Fluidigm) to measure markers of T, B and NK cell lineage, naïve (N) central and effector memory (CM and EM) differentiation, exhaustion, chemokine receptors and transcription factors to characterize cell populations implicated in clinical alloreactivity in prior studies of immune reconstitution after AHST. For a subset of samples, a secondary panel with additional NK cell markers was also employed. Patient peripheral blood mononuclear cells (PBMC) were CD45-barcoded, spiked with CD45-barcoded PBMC from a single healthy control, and acquired on a CyTOF2 mass cytometer. Control cells were used to normalize patient cells using an R code algorithm to minimize batch effects.(25) Further details are in **Tables S1-4** and **Figure S1.**

### High dimensional analysis

The MC data analysis pipeline had four phases. First, cleaning and normalizing data, second visualising data through dimensionality reduction, third partitioning and extracting information using clustering algorithms and finally comparisons between patients grouped by clinical outcomes. FlowJo software (v10) was used to clean data files to viable CD45^+^ single cells. FCS files at each time-point were then resolved, normalised, concatenated, visualized into tSNE plots and clusters using PhenoGraph(26) and FlowSOM(27) algorithms repeated multiple times to ensure stability and robustness. Orthogonal validation of high dimensional clustering data analysis included Boolean determination of population frequencies and use of complimentary supervised learning algorithms including Citrus(28) and Diffcyt(29) via Cytobank and R respectively.

### Frequency distribution, principal components analysis (PCA) and network visualization

Frequency distribution of FlowSOM cell clusters was assigned as normal. skewed or bi-modal by Quantile-Quantile (QQ) plot pattern analysis using SPSS Statistics 25 (IBM).(30) PCA was performed on FlowSOM clusters using Prism v9 (GraphPad). Principal components with the top two Eigenvalues were selected by Scree Plot analysis confirmed by multiple Monte Carlo simulations. Cellular regulatory networks were generated with effector T-cells as target nodes, immunoregulatory cell populations negatively correlated with effector T-cells as source nodes and correlation strength as edges using Cytoscape v3.9.1 (Cytoscape Consortium).

### Cell Sorting and Suppression assays

Samples were sorted with a FACS Aria Fusion cell sorter (Beckton Dickinson) into CD56^hi^ and CD56^hi^CD45RA^pos^ fractions with purity above 90%. Suppression assays with CFSE-labelled patient PBMC responders and irradiated allogeneic PBMC stimulators were performed as previously described.

### Tumor-associated antigen (TAA)-specific T-cell enumeration and measurement of serum biomarkers

CD8 T-cells specific for HLA-A201-restricted peptides from WT-1 and Proteinase 3 TAAs(31) from HLA-A201^+^ patients were enumerated with dextramers (Immudex). Serum biomarkers REG3α and ST2(32) were measured in patient serum using Luminex multiplexing ELISA technology.

### Other statistical considerations

Frequencies of individual immune cell clusters generated in PhenoGraph, FlowSOM and Citrus were compared using unpaired Mann Whitney tests. P values were uncorrected for multiple comparisons. Differential abundance of individual immune cell clusters in Citrus and Diffcyt were compared using p values adjusted for false discovery/multiple comparisons. Cumulative incidence (CI) of aGvHD was calculated up to D+180 to capture both classical and late-onset aGvHD(33), with relapse and treatment-related mortality as competing risks.

Further details of the AHST platform and outcome assessments, MC pipeline and experimental approaches are in **Supplemental Methods.**

## Results

### Study design, sample accrual and analysis pipeline

Peripheral blood samples were prospectively collected at 5 time-points from 58 consecutive patients undergoing RIC-AHST with uniform conditioning and post-transplant immunoprophylaxis. Donor and patient demographics are given in **Table S5.** Samples were processed for MC and at early time-points also for serum protein biomarker analysis and enumeration of TAA-specific T-cells, **Figure S2A**.

Patients were eligible for assays if they remained free of aGvHD and disease relapse. Donor chimerism levels were high at early time-points, with a median CD3 T-cell donor chimerism of 83% (range 47-100%) at D+30. The CI of aGvHD at D+180 was 40%. The median onset of aGvHD was D+92 (range D+31-D+165) with a biphasic incidence over time. Of patients developing aGvHD, half did so by D+60 and half after D+90 during or after reduction of CNI. Five patients relapsed, 15 had clinically significant CMV reactivation and 4 had other significant viral infections during the study. Sample accrual was high with 189 of 232 eligible samples (82%) collected and meeting QC parameters for inclusion in the MC pipeline, **Figures S2B-D and S3**.

### The cellular immune landscape at D+30

Samples from 56 patients at the first time-point after AHST were processed and passed QC for acquisition using the primary MC panel. CD45-barcoded patient cells were easily resolved from spiked healthy controls with a median of 112,362 viable CD45^+^ events acquired. High dimensional data visualization post-normalization as tSNE plots produced defined regions of immune cells expressing different lineage markers, **Figure 1A**. The use of complimentary clustering algorithms, PhenoGraph and FlowSOM, reproducibly identified 24 and 40 phenotypically distinct clusters respectively, **Figure 1B-C**.and validated using Boolean analysis. Frequencies of higher order phenotypic groups (HOPG) were concordant across both clustering models with those generated with FlowSOM clustering most closely correlated to Boolean analysis, **Figure 1D**.

**Figure 1.**
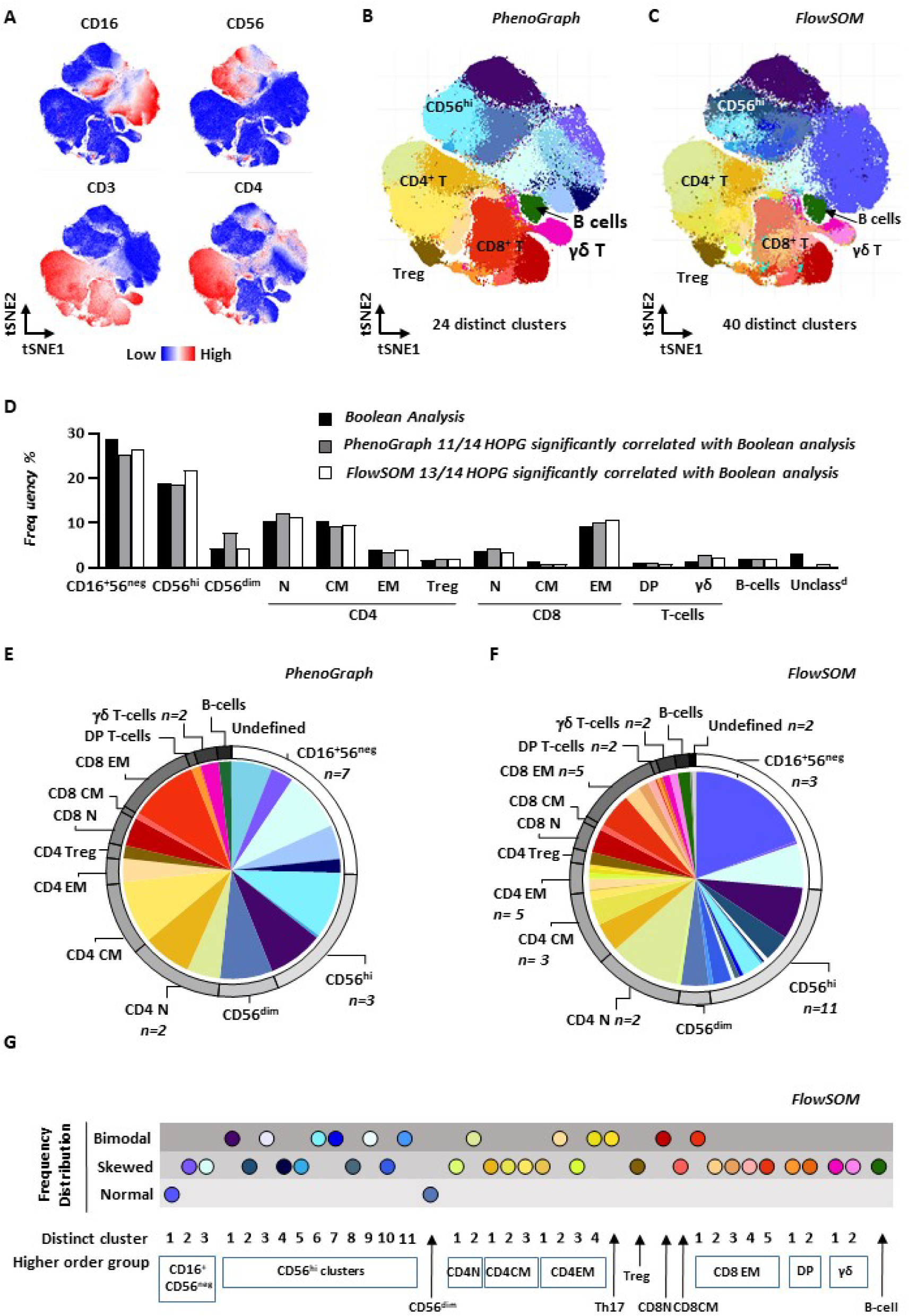
The cellular immune landscape at D+30 after AHST. **A** High dimensional visualization of peripheral blood immune cells. tSNE plots are shown with expression levels of major lineage markers. **B-C** PhenoGraph and FlowSOM clustering algorithms identified 24 and 40 phenotypically distinct T-cell clusters respectively. tSNE plots with individual clusters differentially coloured are shown. **D** Frequencies of higher order phenotypic groups (HOPG) were similar using either PhenoGraph or FlowSOM clustering algorithms or Boolean gating analysis. N naïve; CM, central memory; EM, Effector Memory; Treg, regulatory T-cell; DP, double positive T-cell. **E-F** Pie and arc diagram depicting relative frequencies of phenotypically distinct individual clusters identified with PhenoGraph and FlowSOM expressed as percentage of viable CD45^+^ lineage^+^ cells (slices) and their HOPG (arcs). The models detected distinct clusters with frequencies above 0.3%. **G** Frequency distribution patterns of individual FlowSOM clusters identified by Q-Q plot analysis. Clusters within CD56^hi^ NK cell and CD4 and CD8 EM T-cell HOPGs exhibited bimodal distribution patterns. Data is from 56 individual patient samples.

Both clustering models assigned 98% of cells into discrete clusters within defined HOPGs with a lower limit of detection of frequency of 0.3% of viable CD45^+^ lineage^+^ cells, **Figure 1E-F**. Invariant NKT-cells were not detectable by clustering models and were enumerated using Boolean analysis (median frequency 0.15% of viable CD45^+^ lineage^+^ cells). High frequencies of CD56^hi^ NK cells were identified in both clustering models with greater numbers of phenotypically distinct clusters within this compartment and within the CD4 and CD8 T-cell compartments with FlowSOM. In view of this, and the closer correlation with Boolean analysis, FlowSOM was selected as the primary clustering model for downstream analysis.

Most individual FlowSOM clusters had normal or skewed distribution across all samples at Importantly, individual clusters within the CD56^hi^ NK and CD4 and CD8 EM compartments had bi-modal distribution patterns suggesting their frequencies may segregate with different patient groups or outcomes, **Figure 1G**.

### A distinct immune signature at D+30 identified patients who subsequently develop aGvHD

PCA of FlowSOM cluster frequencies at D+30 demonstrated patients subsequently developing aGvHD had higher PC1 and PC2 values than patients who remained aGvHD-free, **Figure S3A-B.** Individual clusters contributing positively to these PC included CD4 CM and EM and CD8 EM clusters and negative contributors included multiple CD56^hi^ NK cells clusters, GATA3^pos^CD4 EM, and other cells with known immunoregulatory function including Vδ2 γδ- and DP T-cells, **Figure 2A**.

**Figure 2.**
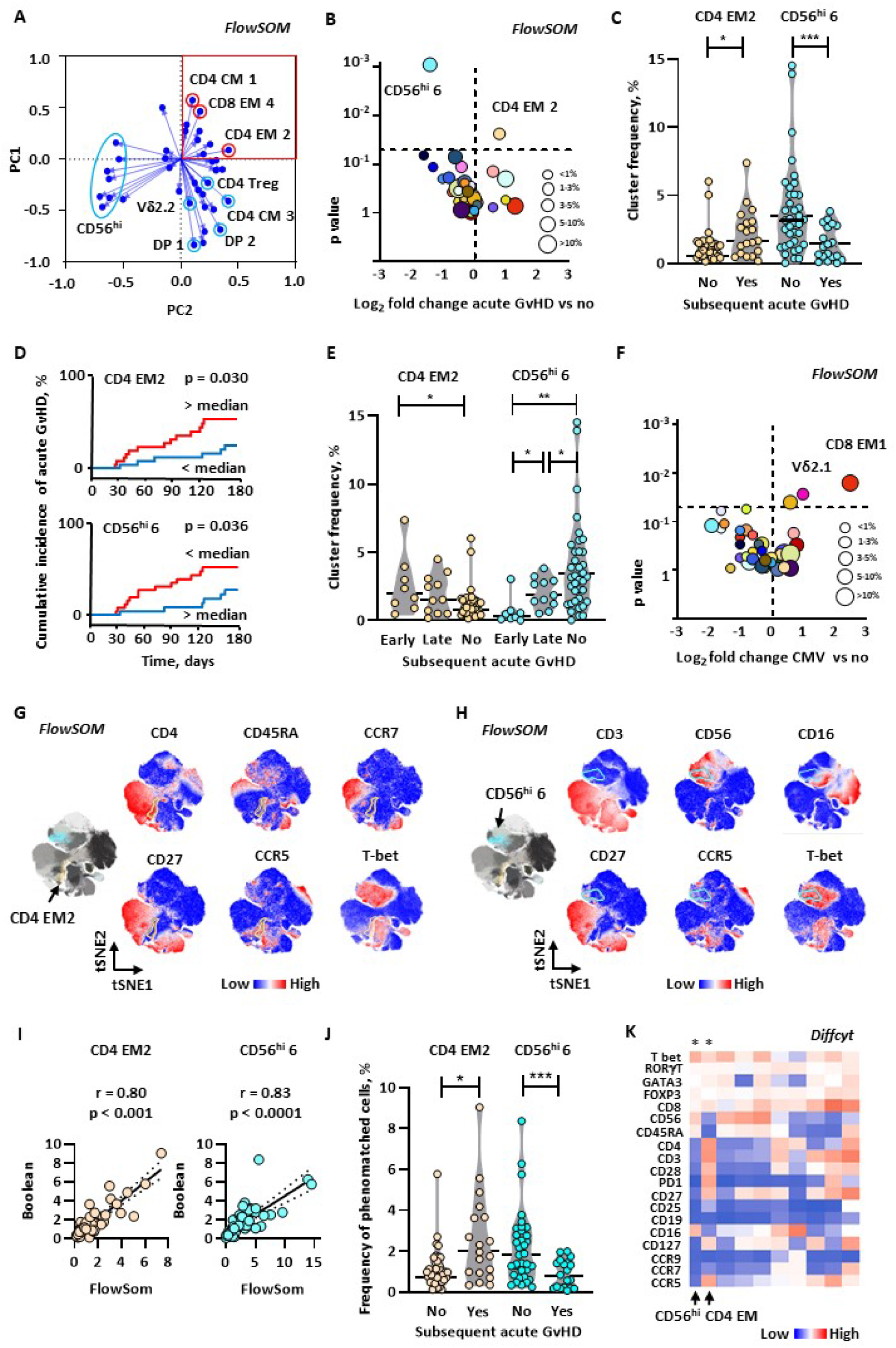
A distinct immune signature at D+30 identified patients who subsequently develop aGvHD. **A** Loadings plot for FlowSOM clusters for the 2 principal components (PC) with largest Eigenvalues. The high PC1 and PC2 region significantly enriched for patients who subsequently developed aGvHD is boxed in red. Individual clusters with the largest positive and negative contributions to the PCs are highlighted with red and blue circles respectively. EM, effector memory; CM, central memory; DP, double positive. **B** Comparison of FlowSOM cluster frequencies identified significantly increased frequencies of CD4 EM cluster 2 and decreased frequencies of CD56^hi^ NK cluster 6 in patients who subsequently developed aGvHD compared to those who did not. Volcano plot depicts fold-change in individual FlowSOM cluster abundance. Data for the whole patient cohort is shown. P values are for Mann Whitney Test (MWT). **C** Frequencies of the two differentially abundant FlowSOM clusters (CD4 EM cluster 2 and CD56^hi^ cluster 6) in individual patients who later developed aGvHDand those who did not. Data for the whole patient cohort is shown. P values are for MWT. **D** Cumulative incidence of aGvHD was significantly greater in patients with CD4 EM cluster 2 frequency above the cohort median and significantly lower in those with CD56^hi^ cluster 6 frequency above the cohort median. P values are for Gray’s Test. **E** Frequencies of the two differential FlowSOM clusters in patients who developed aGvHD early (pre-D+60) or later (post-D+60) compared to those who remained free of aGvHD. Data for the whole patient cohort is shown. P values are for MWT. **F** FlowSOM analysis identified a contrasting immune signature in patients with early CMV reactivation, with increased CD8 EM T-cell and Vδ2.1 γδ T-cell clusters. Volcano plot depicts fold-change in individual FlowSOM cluster abundance in patients who reactivated and cleared CMV prior to D30 compared to those who did not. Data for the whole patient cohort is shown. P values are for Mann Whitney Test (MWT). **G-H** The differential FlowSom CD4 EM cluster 2 had a CD45RA^neg^ CCR7^neg^ CD27^neg^ PD1^neg^ effector memory phenotype with intermediate expression levels of CCR5 and Tbet, whereas the differential CD56^hi^ cluster 6 had a transitional CD16^lo^ CD27^lo^ Tbet^+^ phenotype. tSNE plots depicting individual marker expression levels are shown with the CD4 EM cluster 2 delineated in yellow and the CD56^hi^ cluster 6 delineated in blue. **I-J** Validation of differential FlowSOM clusters using Boolean gating analysis. Frequencies of both differential CD4 EM cluster 2 and CD56^hi^ cluster 6 were closely correlated with frequencies using Boolean analysis to enumerate cells possessing the closest non-redundant phenotype to the FlowSOM clusters (**I**), which were also differentially distributed amongst patients who did/did not subsequently develop aGvHD(**J**). **K** The top 2 differentially abundant clusters in patients who did/did not subsequently develop aGvHD using Diffcyt analysis were CD56^hi^ NK and CD4 EM T-cell clusters, with the same vector of change and similar phenotypes to those identified in FlowSOM clustering and Boolean approaches. *denotes false discovery rate <0.05.

Analysis of individual FlowSOM clusters identified a CD4 EM cluster (CD4 EM cluster 2) and a CD56^hi^ NK cell cluster (CD56^hi^ cluster 6) present at significantly increaased and decreased frequency respectively in patients who subsequently developed aGvHD, **Figure 2B-C**. Patients with unrelated donors had significantly lower frequencies of CD56^hi^ cluster 6 but other pre-transplant factors did not significantly impact frequencies of these clusters, Table S6. The total CD4 EM compartment was not significantly different in the two patient groups nor were frequencies of iNKT-cells enumerated by Boolean analysis. Importantly, the CI of aGvHD was significantly different in patients whose absolute counts of the two differential FlowSOM clusters were above or below median levels for the cohort, **Figure 2D**, but was not significantly impacted by any clinical variable. Differential frequencies of both clusters were more marked in patients who developed aGVHD soon after D+30 **Figure 2E**. In contrast, patients with prior CMV reactivation had a distinct immune signature at D+30 characterized by increased CD8 EM- and γδ-T-cell clusters, both known to contribute to CMV control post-AHST, **Figure 2F**. When these patients were excluded, the immune signature predicting aGvHD was more prominent, Figure S4C. The CD4 EM cluster 2 had a CD27^neg^PD1^neg^ phenotype with intermediate CCR5 and Tbet expression consistent with Th1 alloreactive function, whereas the CD56^hi^ cluster 6 had a transitional CD56^hi^CD16^lo^CD27^lo^Tbet^pos^ phenotype, **Figure 2G-H**.

The Flowsom clustering immune signature associated with subsequent aGvHD was validated across multiple alternate analysis platforms, including Boolean analysis using a best non-redundant phenotype approach, Figures S4D-G and 2I-J, the alternative clustering model PhenoGraph and the supervised learning algorithms Citrus and Diffcyt, where CD4 EM and CD56^hi^ NK cell clusters remained significantly differentially abundant after correction for false discovery rates, Figures S4H-K and 2K.

### CD56^hi^ NK cells phenomatched to FlowSOM CD56^hi^ cluster 6 have a cytolytic phenotype and allosuppressive function

The frequency of FlowSOM CD56^hi^ cluster 6 was significantly negatively correlated with both FlowSOM CD4 EM cluster 2 and donor T-cell chimerism at D+30, consistent with CD56^hi^ cluster 6 cells limiting expansion of alloreactive donor T-cells, **Figure 3A-B**. In view of this finding, we next sought to phenotypically and functionally characterize CD56^hi^ cluster 6 cells We used a secondary MC panel to further characterize CD56^hi^ NK cells in a subset of D+30 patient samples. Utilizing FlowSOM to cluster cells stained with our secondary panel we identified the cluster most closely phenomatching FlowSOM CD56^hi^ cluster 6 identified with our primary panel. The phenomatched cluster had high expression of perforin, but did not express NKG2A, NKp46, intracellular IL-10 or TGFβ, **Figure 3C**.

**Figure 3.**
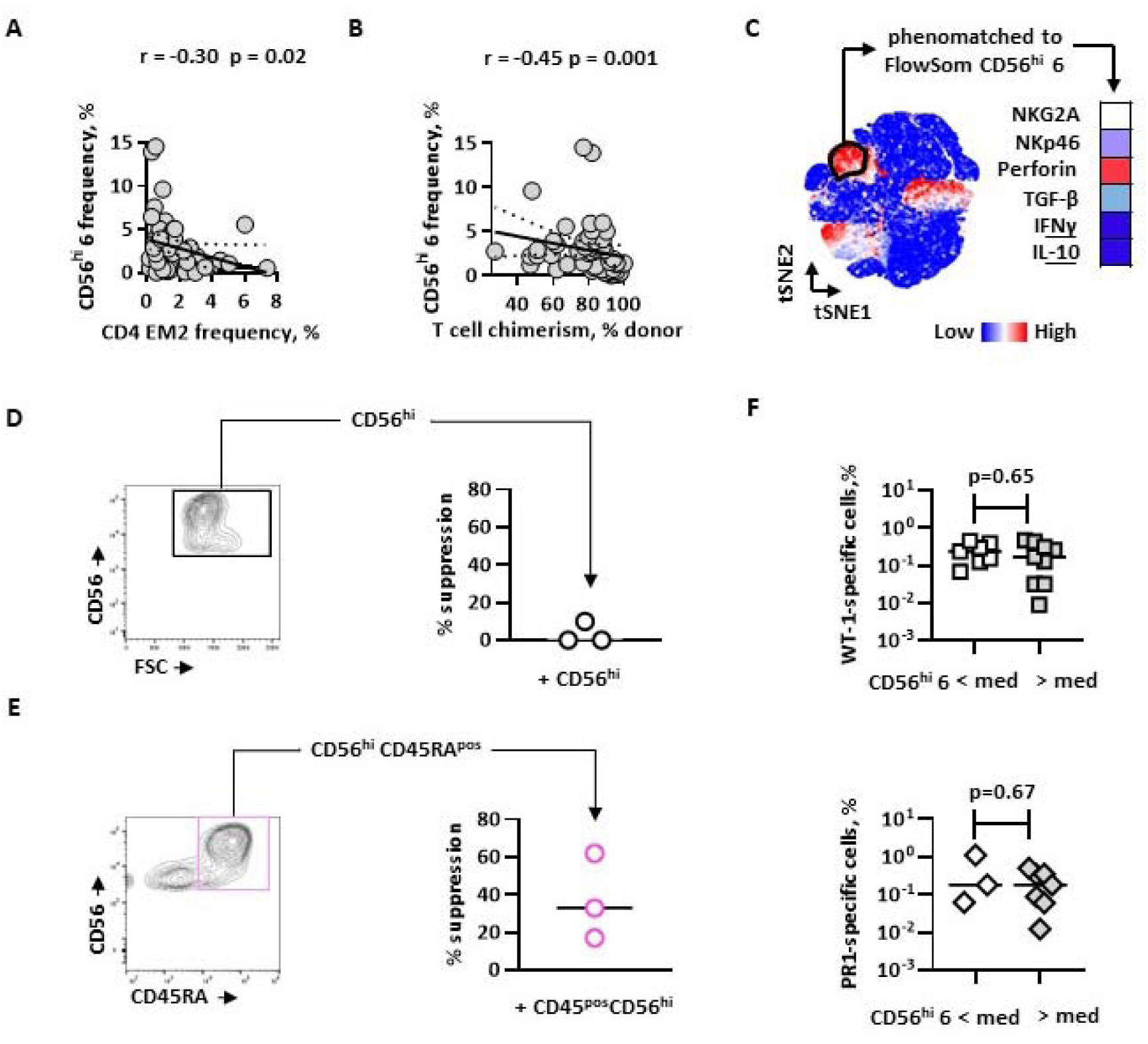
CD56^hi^ NK cells deficient in patients who developed subsequent aGvHD have a cytolytic phenotype and possess functional allosuppressive capacity. **A-B** The frequency of the FlowSOM CD56^hi^ NK cell cluster increased in patients who remained free of aGvHD (FlowSOM CD56^hi^ cluster 6) was negatively correlated with the frequency of the FlowSOM CD4 EM cluster 2 increased in patients who subsequently develop aGvHD, and also with donor CD3 T-cell chimerism levels at D+30. R values are for Spearman correlations. **C** CD56^hi^ cells phenomatched to FlowSOM CD56^hi^ cluster 6 possessed a NKG2A^dim^ NKp46^neg^ Perforin^hi^ phenotype distinct to the whole CD56^hi^ cell pool. Heat map depicts proportion of viable unstimulated CD56^hi^ cells from a cluster phenomatched to FlowSOM CD56^hi^ cluster 6 from patients at D+30 expressing individual molecules determined by single-cell mass cytometry. Results are from 7 patient samples. **D** Whole populations of CD56^hi^ NK cells purified on expression of CD56 alone did not suppress T-cell alloresponses. Results are depicted from 3 separate D+30 patient samples. Horizontal lines are medians. **E** Purified CD45RA^pos^ CD56^hi^ NK cells suppressed T-cell alloresponses. Results are depicted from 3 separate D+30 patient samples. Horizontal lines are medians. **F** Expansion of FlowSOM CD56^hi^ cluster 6 cells was not associated with reduced TAA-specific T-cell frequencies *in vivo*. Results are for 16 HLA-A0201 patient samples at D+30. Horizontal lines are medians. P values are for MWT. Med, median frequency. Frequencies are expressed as percentage of live CD8^+^ T-cells.

To ascertain if CD56^hi^ NK cells deficient in patients subsequently developing aGvHD had capacity to suppress alloreactive T-cell responses we sorted cells from D+30 patient samples and added them to allostimulated patient PBMC. Whole populations of CD56^hi^ NK cells did not suppress T-cell alloproliferation, **Figure 3D**. Flowsom CD56^hi^ cluster 6 did not have a non-redundant surface phenotype to directly sort them from other CD56^hi^ NK cells, **Figure S4F,** so we next purified CD45^pos^ CD56^hi^ NK cells to enrich for cluster 6. Notably, CD45RA^pos^CD56^hi^ NK cells suppressed T-cell alloproliferation, **Figure 3E**, confirming cells partially phenomatched to those increased in patients remaining aGvHD-free have allosuppressive capacity, and demonstrating this functionality is restricted to sub-populations of the CD56^hi^ NK cell compartment.

In view of these findings, we finally sought to determine if increased numbers of FlowSom CD56^hi^ cluster 6 NK cells at D+30 were associated with reduced reconstitution of beneficial TAA-specific T-cell populations *in vivo*. Importantly, CD8 T-cells specific to peptide epitopes of the WT1 and Proteinase 3 were detectable at similar frequencies in patients with FlowSOM CD56^hi^ cluster 6 frequencies above or below median levels, suggesting that the presence these cells was not associated with a high degree of wider immunosuppression, **Figure 3F**.

### Network analysis identifies an immunoregulatory cluster group containing additional phenotypically distinct cell populations

To gain a more comprehensive view of interactions between individual immune cell subsets at D+30 we examined correlation patterns between FlowSOM cluster frequencies to construct cellular networks.(34)

Most positive correlations were between clusters from the same HOPG (*cis*). In contrast most negative correlations were between clusters from different HOPGs (*trans*) consistent with immunoregulatory interactions between functionally distinct cell populations, **Figure 4A**. Next, we examined specific correlations between FlowSOM clusters to ascertain if additional cell populations with known immunoregulatory function were negatively correlated with the CD4 EM cluster 2. We identified 6 additional phenotypically distinct clusters comprising 4 phenotypically heterogeneous CD56^hi^ clusters (CD56^hi^ clusters 2, 4, 7 and 11), a CD4^+^CD8^+^ DP cluster (DP cluster 1) and a γδ T-cell population (Vδ2 cluster 2) that along with CD56^hi^ cluster 6 collectively formed a putative *immunoregulatory cluster group* (IRCG), **Figure 4B-C**. Notably CD4 Treg were not constituents of the IRCG at this early time-point.

**Figure 4.**
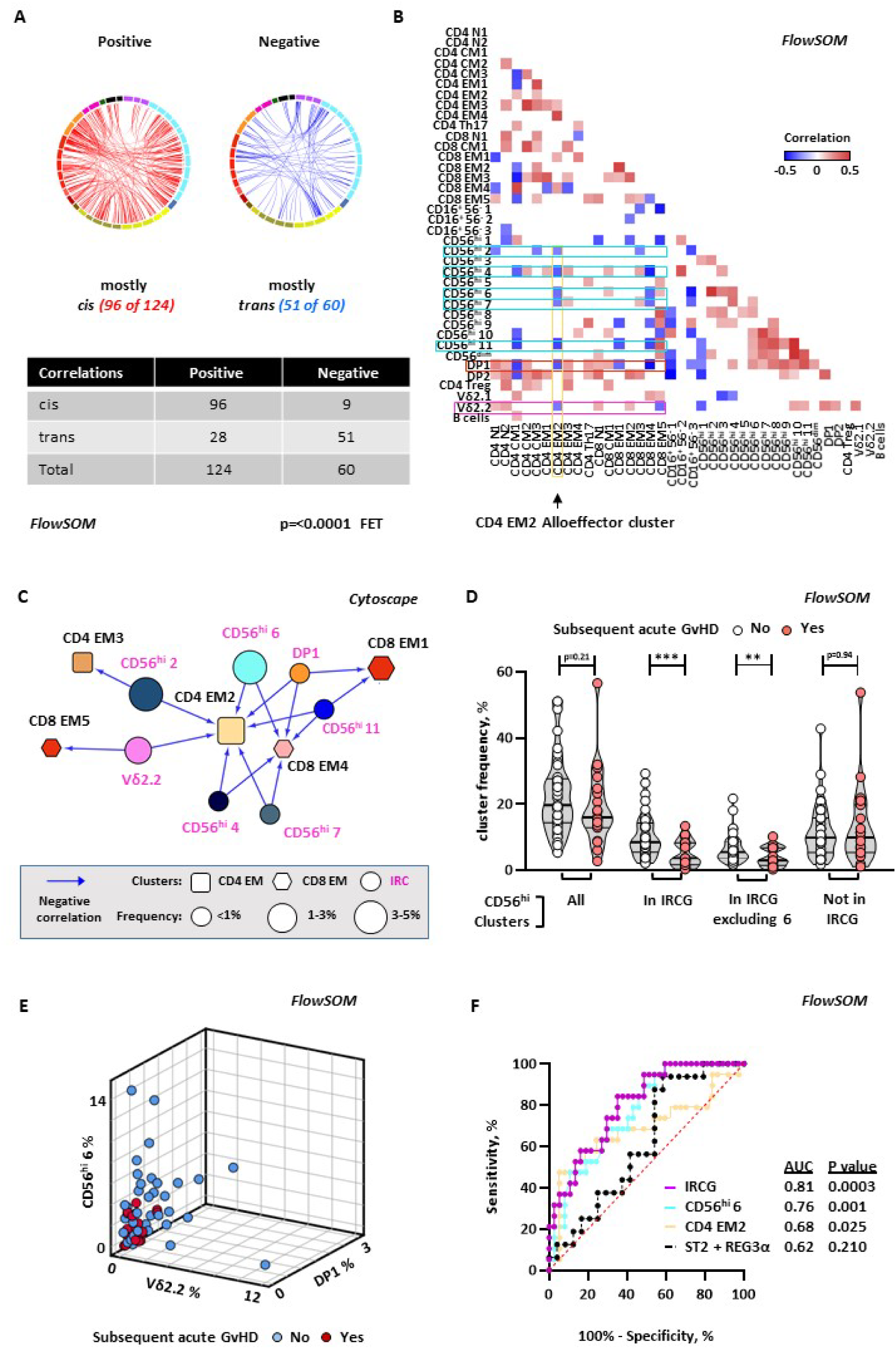
Network analysis identifies an immunoregulatory cluster group containing multiple phenotypically distinct T-cell populations. **A** Positive correlations were mostly between FlowSOM clusters in the same higher order phenotypic groups (HOPG), (*cis* relationships) whereas negative correlations were mostly between clusters in different HOPG (*trans* relationships). Figure depicts chordograms showing positive (red) and negative (blue) correlations between D+30 Flowsom clusters. Only significant (p<0.05) Spearman correlations with r > +/- 0.25 are shown. Embedded table shows numbers of *cis* and *trans* positive and negative correlations. P value is for Fishers Exact Test. **B** Correlation analysis identified FlowSOM clusters (blue boxes) with known immunoregulatory function with significant negative *trans* correlations with the alloreactive CD4 EM cluster 2 (yellow box). Correlation matrix heat map depicts only significant (p<0.05) Spearman correlations with r > -0.30. **C** Cytoscape network depicting the seven phenotypically distinct immunoregulatory clusters significantly negatively correlated with the alloreactive CD4 EM cluster 2, which together form the immununoregulatory cluster group (IRCG). Figure depicts an edge-weighted spring-embedded network; Flowsom clusters are nodes and R values for significant negative Spearman correlations between *trans* clusters are edges (blue arrows) Edge length is inversely proportional to R value. Nodes are scaled to relative cluster frequency. **D** Frequency of Flowsom CD56^hi^ clusters in patients who did and did not subsequently develop aGVHD. P values are for MWT. ** p<0.01, *** p< 0.001. **E** Frequencies of components of the IRCG were not closely correlated with each other. Figure depicts frequencies of CD56^hi^ cluster 6, DP cluster 1 and Vδ2 cluster 2 frequencies in individual patients who did (red) and did not (blue) subsequently develop aGvHD. **F** Receiver-operator curve analysis of the sum of the frequency of clusters within the IRCG, frequencies of individual CD4 EM cluster 2 and CD56^hi^ cluster 6, and ST2 and REG3α serum biomarkers for subsequent development of aGvHD. AUC, area under the curve.

To provide further evidence of the immunoregulatory function of IRCG clusters we compared their frequencies in patients who did and did not subsequently develop aGvHD. Importantly, the summed frequency of CD56^hi^ clusters in the IRCG was significantly lower in patients with subsequent aGvHD, even if CD56^hi^ cluster 6 was excluded, whereas CD56^hi^ clusters not in the IRCG were not reduced, **Figure 4D**. Constituent clusters within the IRCG were not closely correlated, consistent with non-redundancy between different immunoregulatory cell populations, **Figure 4E**. Importantly, the IRCG frequency was a better predictor of subsequent aGvHD than individual CD4 EM cluster 2 or CD56^hi^ cluster 6, or the combination of two validated serum biomarkers (ST2 and REG3α), **Figure 4F**. Taken together these data provide evidence that multiple phenotypically distinct immunoregulatory cells including contribute to the control of CD4 T-cell alloresponses at this early time-point after AHST.

### Alloreactive T-cell signatures preceding aGvHD change over time

To delineate the evolution of cellular immune signatures preceding aGvHD we used clustering algorithms with similar parameters to analyse patient samples from later time-points. HOPG frequencies in clustering models maintained close concordance with Boolean analysis, demonstrating expansion of CD4 and CD8 EM, CD56^dim^ NK cells and B cell populations over time, **Figure S5A-B.**

FlowSOM clustering identified 38-40 clusters at later time-points, of which 95-100% could be assigned a HOPG. New distinct clusters emerged within HOPGs increasing over time whereas the number of phenotypically distinct clusters within HOPGs contracting over time reduced, **Figure 5A-B**.

**Figure 5.**
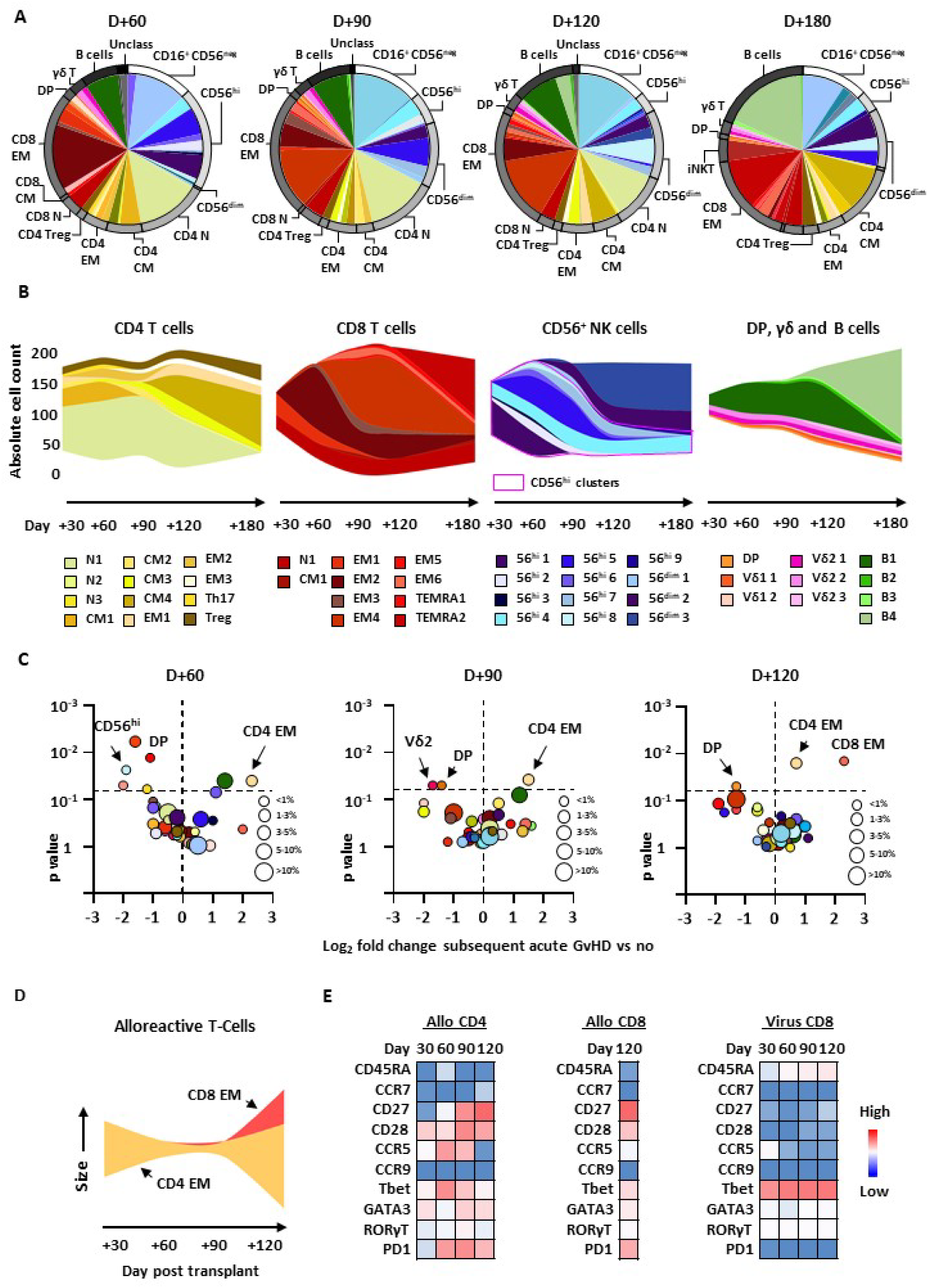
Cluster analysis at later time-points after AHST delineates the evolution of alloreactive T-cell signatures preceding development of aGvHD. **A** Relative frequencies of individual clusters and HOPG identified by Flowsom cluster analysis at later time-points. Diagrams depict relative frequencies of individual clusters (expressed as percentage of viable CD45^+^lineage^+^ cells, slices) and HOPG (arcs). **B** New phenotypically distinct clusters arose in expanding HOPG whereas numbers of phenotypically distinct clusters within contracting HOPG decreased over time. Figures depict stream plots for absolute cell count (cells/µL) of HOPG and constituent clusters over time. Closely phenotypically related clusters are assigned the same colour across different time-points. **C** Allospecific immune signatures at later time-points. Volcano plots depict fold change in individual Flowsom cluster frequencies. To reduce confounding effects of viral immune signatures we previously identified, patients with clinically significant viral reactivation/infection were excluded. P values are for MWT **D** The CD4 EM component of the alloreactive signature was maintained over time, joined later by a CD8 EM component. Figure depicts a stream plot showing relative size of alloreactive CD4 and CD8 compartments over time. **E** Phenotypic evolution of alloreactive EM T-cells over time. Heat maps depict median marker expression level for cells in CD4 and CD8 EM T-cell clusters expanded in patients who subsequently developed aGvHD, and the CD8 EM T-cell cluster expanded in patients with viral reactivation/infection.

To identify predictive immune signatures at later time-points we compared frequencies of FlowSOM clusters in patients grouped by outcome. CD4 EM clusters were significantly increased at all later time-points in patients subsequently developing aGvHD, whereas CD8 EM clusters with increased frequency only became apparent later. The increased T-cell cluster frequencies predictive of subsequent aGvHD at later time points were accompanied by reduced frequencies of several distinct immunoregulatory clusters. **Figure 5C-D**. In contrast, FlowSOM immune signatures associated with viral reactivation/infection contained expanded CD8 EM clusters at all later time-points, Figure S5C FlowSOM immune signatures predictive of aGvHD at given time-points showed intermediate similarity with aGvHD signatures at preceding time-points, but remained dissimilar to viral signatures at each time-point, Figure S5D. Importantly, the same vector and similar magnitudes of change were also seen in analogous cell populations in patients grouped by subsequent aGvHD or viral reactivation/infection using PhenoGraph clustering, Figure S5E

The phenotype of FlowSOM CD4 EM clusters associated with subsequent aGvHD changed over time with sustained expression of CD28, a peak in CCR5, Tbet and PD1 expression followed by a later increase in CD27 expression. The phenotype of CD8 EM expanded at D+120 in patients subsequently developing aGvHD was distinct from CD8 EM in the viral signature, which remained relatively stable over time, **Figure 5E**. Taken together these findings demonstrate phenotypic evolution over time of the alloreactive T-cell compartment preceding aGvHD.

### The IRCG associated with freedom from aGvHD also evolves over time

We systematically analysed correlation patterns of phenotypically distinct FlowSOM clusters to construct networks of EM T-cells and immunoregulatory cell populations at later time-points after AHST. Applying the same criteria used in our D+30 analysis we identified clusters with known immunoregulatory function significantly negatively correlated with CD4 and CD8 EM clusters forming the IRCG at later time-points. **Figure S6A.** The number of phenotypically distinct clusters within the IRCG declined over time, and target clusters shifted from predominantly CD4 to CD8 EM clusters, **Figure 6A**.

**Figure 6.**
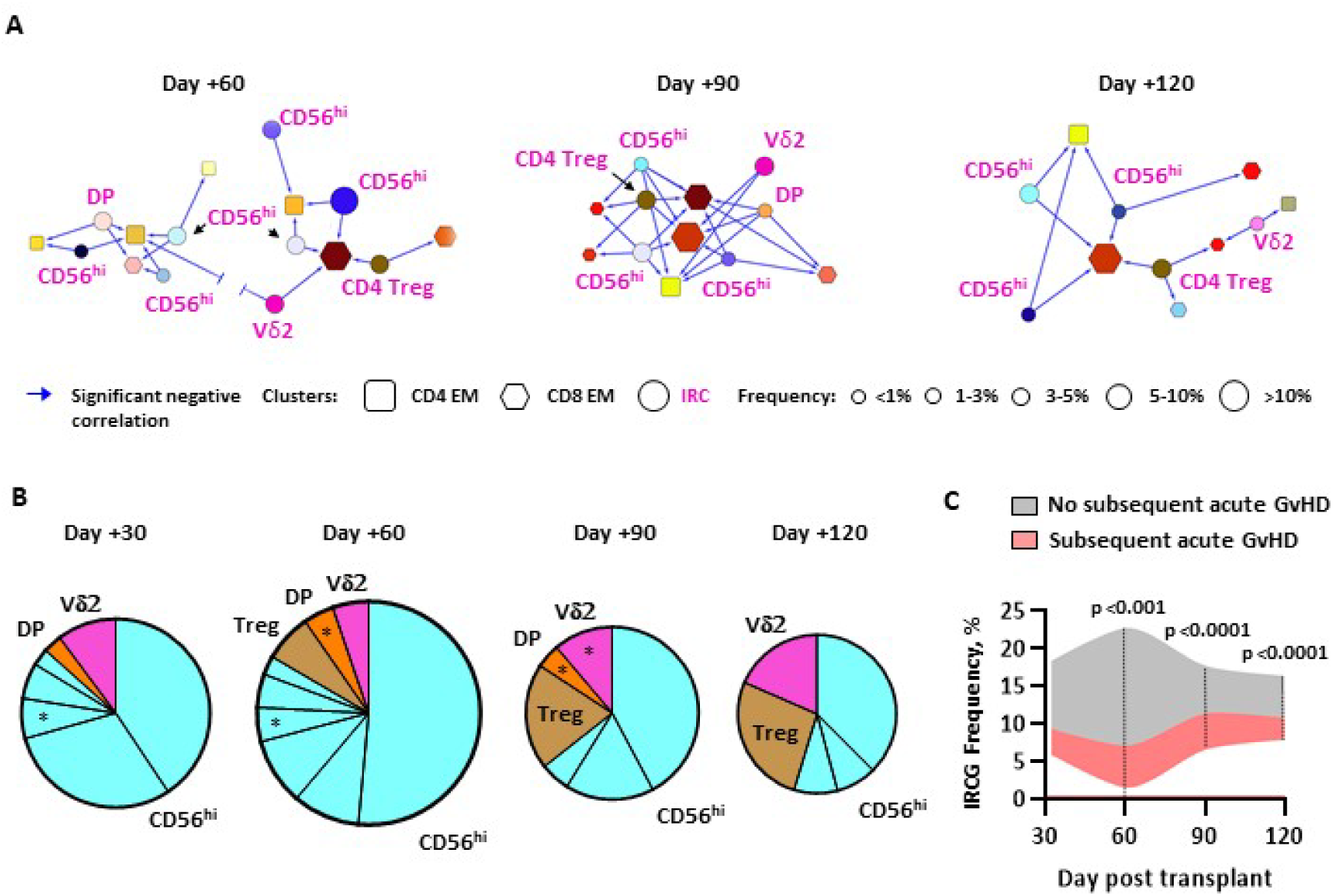
The immunoregulatory cluster group associated with freedom from aGvHD also evolves over time. **A** Cytoscape networks generated from FlowSom clustering models depicting phenotypically distinct immunoregulatory clusters significantly negatively correlated with CD4 and CD8 EM clusters at later time points. Figures depict edge-weighted spring-embedded networks; clusters are nodes and R values for significant negative Spearman correlations between *trans* clusters are edges (blue arrows). Edge length is inversely proportional to R value. Nodes are scaled to relative cluster frequency. **B** Constituent clusters forming the IRCG change over time after AHST. Pie charts are shown depicting the relative proportions of IRCG component clusters at each time-point. Pie size is proportional to the IRCG frequency, which increased and then contracted over time. * denotes individual clusters that with significantly higher frequencies in patients who remained free of aGvHD. **C** Frequency of IRCG over time in patients who did (pink) and did not (grey) subsequently develop aGvHD. Median values across the whole cohort are shown at each time-point. P values are for MWT for AUC at each time-point, *** p< 0.001, **** p< 0.0001

The size of the IRCG and its constituent clusters also changed over time. The proportion of CD56^hi^ clusters reduced whereas CD4 Treg formed an increasing proportion of the IRCG form D+60 onwards. In contrast, DP T-cells were a constituent of the IRCG until D+90 and Vδ2 γδ T-cells were throughout, **Figure 6B**.

To determine if the association with freedom from subsequent aGvHD was maintained at later time-points we performed an area-under-the curve (AUC) analysis of IRCG frequencies over time. Importantly, the AUC of IRCG frequency at later time-points was significantly greater in patients who remained free of aGvHD, **Figure 6C**. Over time, the largest CD56^hi^ clusters within the IRCG maintained a transitional NK phenotype but exhibited increased expression of CD27. DP T-cells maintained a CD8^weak^ phenotype and Vδ2 γδ T-cells and CD4 Treg within the IRCG remained phenotypically stable, **Figure S6B**

Together these data identify changes over time in individual immunoregulatory cell populations contributing to the control of T-cell responses after AHST, with potential significance for design of strategies prevent or reduce aGvHD.

## Discussion

In this study we used single-cell MC to characterize immune cells in patient peripheral blood after AHST. Early studies using this technology reported small heterogeneous patient groups focusing on immune signatures at or after onset of aGvHD. In contrast, we studied a larger prospective patient cohort with a uniform AHST platform to identify a robust early immune signature predictive of subsequent aGvHD. and show how the cellular components within immune signatures predictive of aGvHD evolve over time.

The early immune signature we identified predicting subsequent aGvHD contained increased CD4 EM cells. Previous studies have shown human alloreactive CD4 T-cells in donor cell pools lie mainly within naïve and CM compartments.(6, 35) CD4 EM cells expand i*n vivo* after AHST,(36) and increased numbers of peripheral blood CD4 EM cells have been reported prior to aGvHD(37) consistent with differentiation of alloreactive CD4 cells after encountering alloantigen. In line with this, the CD4 EM cells in our early immune signature had a CD27^neg^ *EM4* phenotype(38) potentially reflecting downregulation of CD27 occuring with sustained antigenic stimulation.(39)

In contrast, CD8 T-cells were not a component of the early aGvHD immune signature even though most minor histocompatibility antigens (mHag) eliciting alloreactive T-cell responses in HLA-matched AHST are MHC Class I-restricted. However, MHC Class 2-restricted mHag can stimulate CD4 T-cell alloresponses.(40) Some prior studies demonstrated increased CD8 EM cells shortly before or at the onset of aGvHD,(8) but such studies were largely restricted to full-intensity conditioning which creates pro-inflammatory environments. After RIC-AHST alloreactive T-cell expansion is more likely to require additional positive costimulatory signals. Notably, the CD4 EM cluster we identified at early time-points expressed the co-stimulatory molecule CD28, which could provide additional proliferative signals in vivo, whereas CD8 EM clusters were largely CD28^neg^. Alternatively our findings may reflect increased sensitivity of CD8 alloresponses to CNI(41) or early partitioning of these cells into tissues.(42)

The other component of the early immune signature predicting subsequent aGvHD was deficient numbers of CD56^hi^ NK cells. These findings extend previous observational studies reporting increased GvHD in patients with low graft CD56^hi^ content(15) and low peripheral blood CD56^hi^ NK cells at onset of aGvHD.(14) The CD56^hi^ NK cells in our early immune signature had an intermediate maturation but highly lytic phenotype in contrast to CD56^hi^ NK cells in healthy donors or grafts, but consistent with expanded populations of peripheral blood CD56^hi^ cells seen early after AHST,(43) and expressed high levels of perforin but low levels of basal IL-10 and TGFβ secretion, supporting a cytolytic mechanism of suppression of alloresponses. Furthermore, we demonstrate functional allosuppressive capacity is restricted to CD45RA^pos^ subsets of CD56^hi^ NK cells early after AHST, which could inform future strategies to use these cells to control harmful alloresponses.

Network analysis identified several additional cell populations with immunoregulatory capacity forming the IRCG that was more accurately predictive of subsequent aGvHD than any individual cell subset identified by differential distribution analysis. At the earliest time-point, the IRCG included additional sub-populations of CD56^hi^ NK cells suggesting that control of CD4 T-cell alloresponses is not restricted to a single phenotypic group of these cells. Additionally, the IRCG contained Vδ2 γδ T-cells and DP T-cells. Although early clinical studies linked higher γδ T-cell counts and increased aGVHD,(44) subsequent studies showed the Vδ2 γδ T-cell subset was associated with less aGvHD,(18) and Vδ2 cells can suppress CD4 alloproliferation *in vitro.*(45) Notably, the DP T-cell cluster within the IRCG was CD8^weak^ consistent with a CD8^α/α+^ immunoregulatory phenotype rather than a CD8^α/β+^ pro-inflammatory phenotype.(46)

Immune signatures associated with subsequent aGvHD changed over time. Although alloreactive CD4 EM cells remained a constant component, these cells exhibited 3 phenotypic phases-an early *EM4* CCR5^pos^Tbet^pos^PD1^neg^ phenotype indicating retention of migratory capacity to secondary lymphoid organs and the absence of exhaustion, a mid-phase with maximal Tbet and increasing PD1 expression, followed by a late-phase reduction in Tbet and CCR5 expression and emergence of a CD27^pos^ *EM1* phenotype. This is unlikely to represent phenotypic evolution of a single cell population as CD27 re-expression on effector T-cells rarely occurs, but more likely represents emergence of a transitional *CM/EM1* alloreactive cell population upon reduction of CNI, known to suppress CD27 upregulation(47) and NF-κB-signalling important to CD27-mediated proliferation/survival signals.(48) The emergence of a population of CD27^pos^CD4 T-cells was accompanied by a CD27^pos^CD8 *EM1* T-cell population. Importantly, these findings are consistent with phenotypically distinct alloreactive T-cell populations contributing to development of early and late-onset aGvHD after RIC-AHST.

CNI withdrawal may also provide one mechanism underlying the change in components of the IRCG over time. CD56^hi^ NK cells, also relatively resistant to CNI,(49) were the dominant IRCG component at early time-points but these populations contracted later after AHST concurrent with withdrawal of CNI. Similarly, CD4 Treg, which are known to be more sensitive to CNI than effector T-cells because of their dependence on IL-2 for proliferation,(50, 51) became an increasingly prominent IRCG component at later time-points. Alternatively the emergence of CD4 Treg as an IRCG component may represent the development of alloantigen-specific inducible Treg diverted from the CD4 effector pool as *in vivo* tolerance develops.

Our results support the time-dependent refinement of strategies to target both alloreactive T cells and immunoregulatory cell populations to reduce aGvHD after RIC-AHST. Specifically, the evolving phenotype of alloreactive T-cells suggests strategies targeting the CD27 pathway may be less effective at controlling aGvHD at early time points than limiting the development of late onset aGvHD, Similarly, the early prominence of CD45RA^pos^CD56^hi^ NK cells provides an alternative or additional immunoregulatory cell population to conventional CD4 Treg to target to reduce aGvHD early after AHST.

We used vigorous quality control and cross-validation with multiple algorithms to increase the robustness of our findings. However, there are some limitations to our study. We focused on the T-, B- and NK cell compartments and did not enumerate myeloid cell subsets, which have also been implicated in the control of T-cell alloresponses. Although advantages of our study include the use of a single AHST platform, conditioning regimen, post-transplant immunoprophylaxis and prospective sample acquisition, further work will need to extend our findings to other AHST platforms employing antibody-mediated T-cell depletion or post-transplant cyclophosphamide. Finally, our study was not designed or powered to identify immune signatures predictive of cancer relapse. However, as the alloreactive T cell response also contributes to immune graft-versus-tumor effects, further studies will be needed to delineate the relationship between the immune signatures we have identified, particularly IRCG frequency, and the risk of subsequent cancer recurrence.

In conclusion, using MC we identified immune cell signatures at different times after RIC-AHST associated with subsequent development of aGvHD, providing mechanistic insight into the *in vivo* evolution of human T-cell alloresponses in this context. Furthermore, we demonstrate that phenotypically distinct immunoregulatory cell populations make different contributions to the control of alloresponses at different times. These findings support the development of strategies to target different immune cell populations at different times to prevent or control aGvHD.

## Acknowledgements

We thank Sofie Van Gassen (VIB Inflammation Research Centre, Ghent University, Belgium) for providing the code for the normalization algorithm prior to its commercial availability.

## Authorship Contribution

JA developed and validated the MC pipeline, acquired and analyzed patient peripheral blood samples and performed cell sorting and allogeneic co-culture experiments, and wrote the manuscript. SC assisted with experimental design, cell sorting and allogeneic co-culture experiments and data analysis. ST and MM contributed to experimental design and/or data analysis. SVG provided the code for the normalization process. WJL, MEC and DP supplied reagents and expertise for enumeration of γδ T-cells. JC contributed patient samples, and to the writing of the manuscript, JGG contributed to the conception of the project, patient samples, experimental design and the writing of the manuscript. JKD conceived, designed and supervised the study, analyzed and interpreted experiments and clinical data and wrote the manuscript. All authors approved the final manuscript.

## Data Availability Statement

MC FCS files will be placed in flow repository.org upon publication

For data not publicly accessible contact the corresponding author.

## References

1. Anasetti C, Logan BR, Lee SJ, Waller EK, Weisdorf DJ, Wingard JR, et al. Peripheral-blood stem cells versus bone marrow from unrelated donors. The New England journal of medicine. 2012;367(16):1487–96.

2. Saber W, Opie S, Rizzo JD, Zhang MJ, Horowitz MM, Schriber J. Outcomes after matched unrelated donor versus identical sibling hematopoietic cell transplantation in adults with acute myelogenous leukemia. Blood. 2012;119(17):3908–16.

3. Wagner JE, Thompson JS, Carter SL, Kernan NA, Unrelated Donor Marrow Transplantation T. Effect of graft-versus-host disease prophylaxis on 3-year disease-free survival in recipients of unrelated donor bone marrow (T-cell Depletion Trial): a multi-centre, randomised phase II-III trial. Lancet. 2005;366(9487):733–41.

4. Sengsayadeth S, Savani BN, Blaise D, Malard F, Nagler A, Mohty M. Reduced intensity conditioning allogeneic hematopoietic cell transplantation for adult acute myeloid leukemia in complete remission - a review from the Acute Leukemia Working Party of the EBMT. Haematologica. 2015;100(7):859–69.

5. McCurdy SR, Radojcic V, Tsai HL, Vulic A, Thompson E, Ivcevic S, et al. Signatures of GVHD and relapse after posttransplant cyclophosphamide revealed by immune profiling and machine learning. Blood. 2022;139(4):608–23.

6. Yakoub-Agha I, Saule P, Depil S, Micol JB, Grutzmacher C, Boulanger-Villard F, et al. A high proportion of donor CD4+ T cells expressing the lymph node-homing chemokine receptor CCR7 increases incidence and severity of acute graft-versus-host disease in patients undergoing allogeneic stem cell transplantation for hematological malignancy. Leukemia. 2006;20(9):1557–65.

7. Loschi M, Porcher R, Peffault de Latour R, Vanneaux V, Robin M, Xhaard A, et al. High number of memory t cells is associated with higher risk of acute graft-versus-host disease after allogeneic stem cell transplantation. Biology of blood and marrow transplantation : journal of the American Society for Blood and Marrow Transplantation. 2015;21(3):569–74.

8. Khandelwal P, Lane A, Chaturvedi V, Owsley E, Davies SM, Marmer D, et al. Peripheral Blood CD38 Bright CD8+ Effector Memory T Cells Predict Acute Graft-versus-Host Disease. Biology of blood and marrow transplantation : journal of the American Society for Blood and Marrow Transplantation. 2015;21(7):1215–22.

9. Miura Y, Thoburn CJ, Bright EC, Phelps ML, Shin T, Matsui EC, et al. Association of Foxp3 regulatory gene expression with graft-versus-host disease. Blood. 2004;104(7):2187–93.

10. Rieger K, Loddenkemper C, Maul J, Fietz T, Wolff D, Terpe H, et al. Mucosal FOXP3+ regulatory T cells are numerically deficient in acute and chronic GvHD. Blood. 2006;107(4):1717–23.

11. Rezvani K, Mielke S, Ahmadzadeh M, Kilical Y, Savani BN, Zeilah J, et al. High donor FOXP3-positive regulatory T-cell (Treg) content is associated with a low risk of GVHD following HLA-matched allogeneic SCT. Blood. 2006;108(4):1291–7.

12. Alhaj Hussen K, Michonneau D, Biajoux V, Keita S, Dubouchet L, Nelson E, et al. CD4(+)CD8(+) T-Lymphocytes in Xenogeneic and Human Graft-versus-Host Disease. Frontiers in immunology. 2020;11:579776.

13. Eljaafari A, Yuruker O, Ferrand C, Farre A, Addey C, Tartelin ML, et al. Isolation of human CD4/CD8 double-positive, graft-versus-host disease-protective, minor histocompatibility antigen-specific regulatory T cells and of a novel HLA-DR7-restricted HY-specific CD4 clone. Journal of immunology (Baltimore, Md : 1950). 2013;190(1):184–94.

14. Ullrich E, Salzmann-Manrique E, Bakhtiar S, Bremm M, Gerstner S, Herrmann E, et al. Relation between Acute GVHD and NK Cell Subset Reconstitution Following Allogeneic Stem Cell Transplantation. Frontiers in immunology. 2016;7:595.

15. Kariminia A, Ivison S, Ng B, Rozmus J, Sung S, Varshney A, et al. CD56(bright) natural killer regulatory cells in filgrastim primed donor blood or marrow products regulate chronic graft-versus-host disease: the Canadian Blood and Marrow Transplant Group randomized 0601 study results. Haematologica. 2017;102(11):1936–46.

16. Rubio MT, Moreira-Teixeira L, Bachy E, Bouillie M, Milpied P, Coman T, et al. Early posttransplantation donor-derived invariant natural killer T-cell recovery predicts the occurrence of acute graft-versus-host disease and overall survival. Blood. 2012;120(10):2144–54.

17. Chaidos A, Patterson S, Szydlo R, Chaudhry MS, Dazzi F, Kanfer E, et al. Graft invariant natural killer T-cell dose predicts risk of acute graft-versus-host disease in allogeneic hematopoietic stem cell transplantation. Blood. 2012;119(21):5030–6.

18. Minculescu L, Marquart HV, Ryder LP, Andersen NS, Schjoedt I, Friis LS, et al. Improved Overall Survival, Relapse-Free-Survival, and Less Graft-vs.-Host-Disease in Patients With High Immune Reconstitution of TCR Gamma Delta Cells 2 Months After Allogeneic Stem Cell Transplantation. Frontiers in immunology. 2019;10:1997.

19. Bendall SC, Simonds EF, Qiu P, Amir el AD, Krutzik PO, Finck R, et al. Single-cell mass cytometry of differential immune and drug responses across a human hematopoietic continuum. Science. 2011;332(6030):687–96.

20. Lakshmikanth T, Olin A, Chen Y, Mikes J, Fredlund E, Remberger M, et al. Mass Cytometry and Topological Data Analysis Reveal Immune Parameters Associated with Complications after Allogeneic Stem Cell Transplantation. Cell reports. 2017;20(9):2238–50.

21. Hartmann FJ, Babdor J, Gherardini PF, Amir ED, Jones K, Sahaf B, et al. Comprehensive Immune Monitoring of Clinical Trials to Advance Human Immunotherapy. Cell reports. 2019;28(3):819–31 e4.

22. van Halteren AGS, Suwandi J, Tuit S, Borst J, Laban S, Tsonaka R, et al. A Unique Immune Signature in Blood Separates Therapy-Refractory from Therapy-Responsive Acute Graft-Versus-Host Disease. Blood. 2022.

23. Davies JK, Taussig DC, Oakervee H, Davies AJ, Agrawal SG, Gribben JG, et al. Long-term follow-up after reduced-intensity conditioning allogeneic transplantation for acute myeloid leukemia/myelodysplastic syndrome: late CNS relapses despite graft-versus-host disease. Journal of clinical oncology : official journal of the American Society of Clinical Oncology. 2006;24(14):e23–5.

24. Davies JK, Taussig D, Oakervee H, Smith M, Agrawal S, Cavenagh JD, et al. Long-term survival with low toxicity after allogeneic transplantation for acute myeloid leukaemia and myelodysplasia using non-myeloablative conditioning without T cell depletion. British journal of haematology. 2013;162(4):525–9.

25. Van Gassen S, Gaudilliere B, Angst MS, Saeys Y, Aghaeepour N. CytoNorm: A Normalization Algorithm for Cytometry Data. Cytometry A. 2020;97(3):268–78.

26. Levine JH, Simonds EF, Bendall SC, Davis KL, Amir el AD, Tadmor MD, et al. Data-Driven Phenotypic Dissection of AML Reveals Progenitor-like Cells that Correlate with Prognosis. Cell. 2015;162(1):184–97.

27. Van Gassen S, Callebaut B, Van Helden MJ, Lambrecht BN, Demeester P, Dhaene T, et al. FlowSOM: Using self-organizing maps for visualization and interpretation of cytometry data. Cytometry A. 2015;87(7):636–45.

28. Bruggner RV, Bodenmiller B, Dill DL, Tibshirani RJ, Nolan GP. Automated identification of stratifying signatures in cellular subpopulations. Proceedings of the National Academy of Sciences of the United States of America. 2014;111(26):E2770–7.

29. Weber LM, Nowicka M, Soneson C, Robinson MD. diffcyt: Differential discovery in high-dimensional cytometry via high-resolution clustering. Commun Biol. 2019;2:183.

30. Loy A, Follett L, Hofmann H. Variations of Q-Q Plots: The Power of Our Eyes! Am Stat. 2016;70(2):202–14.

31. Scheibenbogen C, Letsch A, Thiel E, Schmittel A, Mailaender V, Baerwolf S, et al. CD8 T-cell responses to Wilms tumor gene product WT1 and proteinase 3 in patients with acute myeloid leukemia. Blood. 2002;100(6):2132–7.

32. Hartwell MJ, Ozbek U, Holler E, Renteria AS, Major-Monfried H, Reddy P, et al. An early-biomarker algorithm predicts lethal graft-versus-host disease and survival. JCI insight. 2018;3(16).

33. Filipovich AH, Weisdorf D, Pavletic S, Socie G, Wingard JR, Lee SJ, et al. National Institutes of Health consensus development project on criteria for clinical trials in chronic graft-versus-host disease: I. Diagnosis and staging working group report. Biology of blood and marrow transplantation : journal of the American Society for Blood and Marrow Transplantation. 2005;11(12):945–56.

34. Fonseca Dos Reis E, Viney M, Masuda N. Network analysis of the immune state of mice. Sci Rep. 2021;11(1):4306.

35. Foster AE, Marangolo M, Sartor MM, Alexander SI, Hu M, Bradstock KF, et al. Human CD62L-memory T cells are less responsive to alloantigen stimulation than CD62L+ naive T cells: potential for adoptive immunotherapy and allodepletion. Blood. 2004;104(8):2403–9.

36. Latis E, Michonneau D, Leloup C, Varet H, Peffault de Latour R, Consortium C, et al. Cellular and molecular profiling of T-cell subsets at the onset of human acute GVHD. Blood advances. 2020;4(16):3927–42.

37. Matthews K, Lim Z, Afzali B, Pearce L, Abdallah A, Kordasti S, et al. Imbalance of effector and regulatory CD4 T cells is associated with graft-versus-host disease after hematopoietic stem cell transplantation using a reduced intensity conditioning regimen and alemtuzumab. Haematologica. 2009;94(7):956–66.

38. Koch S, Larbi A, Derhovanessian E, Ozcelik D, Naumova E, Pawelec G. Multiparameter flow cytometric analysis of CD4 and CD8 T cell subsets in young and old people. Immun Ageing. 2008;5:6.

39. Hintzen RQ, de Jong R, Lens SM, Brouwer M, Baars P, van Lier RA. Regulation of CD27 expression on subsets of mature T-lymphocytes. Journal of immunology (Baltimore, Md : 1950). 1993;151(5):2426–35.

40. van Balen P, van Bergen CAM, van Luxemburg-Heijs SAP, de Klerk W, van Egmond EHM, Veld SAJ, et al. CD4 Donor Lymphocyte Infusion Can Cause Conversion of Chimerism Without GVHD by Inducing Immune Responses Targeting Minor Histocompatibility Antigens in HLA Class II. Frontiers in immunology. 2018;9:3016.

41. Auchincloss H, Jr., Winn HJ. Murine CD8+ T cell helper function is particularly sensitive to cyclosporine suppression in vivo. Journal of immunology (Baltimore, Md : 1950). 1989;143(12):3940–3.

42. Santos ESP, Cire S, Conlan T, Jardine L, Tkacz C, Ferrer IR, et al. Peripheral tissues reprogram CD8+ T cells for pathogenicity during graft-versus-host disease. JCI insight. 2018;3(5).

43. Dulphy N, Haas P, Busson M, Belhadj S, Peffault de Latour R, Robin M, et al. An unusual CD56(bright) CD16(low) NK cell subset dominates the early posttransplant period following HLA-matched hematopoietic stem cell transplantation. Journal of immunology (Baltimore, Md : 1950). 2008;181(3):2227–37.

44. Viale M, Ferrini S, Bacigalupo A. TCR gamma/delta positive lymphocytes after allogeneic bone marrow transplantation. Bone marrow transplantation. 1992;10(3):249–53.

45. Traxlmayr MW, Wesch D, Dohnal AM, Funovics P, Fischer MB, Kabelitz D, et al. Immune suppression by gammadelta T-cells as a potential regulatory mechanism after cancer vaccination with IL-12 secreting dendritic cells. J Immunother. 2010;33(1):40–52.

46. Sarrabayrouse G, Bossard C, Chauvin JM, Jarry A, Meurette G, Quevrain E, et al. CD4CD8alphaalpha lymphocytes, a novel human regulatory T cell subset induced by colonic bacteria and deficient in patients with inflammatory bowel disease. PLoS Biol. 2014;12(4):e1001833.

47. Martorell J, Martinez-Caceres E, Rojo I, Vives J. Inhibition of CD27 expression by cyclosporine A; role of IL-2. Transplant Proc. 1992;24(1):125.

48. Yamamoto H, Kishimoto T, Minamoto S. NF-kappaB activation in CD27 signaling: involvement of TNF receptor-associated factors in its signaling and identification of functional region of CD27. Journal of immunology (Baltimore, Md : 1950). 1998;161(9):4753–9.

49. Wang H, Grzywacz B, Sukovich D, McCullar V, Cao Q, Lee AB, et al. The unexpected effect of cyclosporin A on CD56+CD16- and CD56+CD16+ natural killer cell subpopulations. Blood. 2007;110(5):1530–9.

50. Matsuoka K, Koreth J, Kim HT, Bascug G, McDonough S, Kawano Y, et al. Low-dose interleukin-2 therapy restores regulatory T cell homeostasis in patients with chronic graft-versus-host disease. Science translational medicine. 2013;5(179):179ra43.

51. Mantel PY, Ouaked N, Ruckert B, Karagiannidis C, Welz R, Blaser K, et al. Molecular mechanisms underlying FOXP3 induction in human T cells. Journal of immunology (Baltimore, Md : 1950). 2006;176(6):3593–602.

