## Supplementary Materials for "Single-cell mass cytometry maps the evolution of immune cell signatures predictive of acute graft-versus-host disease"

**List of Supplementary Material**

**Supplementary Methods**

**Supplementary Tables S1-6**

**Supplementary Figures S1-6**

**Supplementary Methods**

**Reporting Standards**

The results of this study are reported to the following MIBBI standards; Minimum Information About T-cell assays (MIATA) and Minimum Information about Flow Cytometry (MIFlowCyt) as per <https://fairsharing.org/collection/MIBBI>

**Conditioning, stem-cell source, post-transplant GVHD prophylaxis, supportive care and chimerism determination**

The study recruited consecutive patients undergoing RIC-AHST at our centre between 2016 and 2019. All patients received reduced intensity conditioning with Fludarabine (25mg/m^2^ daily for 5 days) and an alkylating agent (cyclophosphamide 1g/m^2^ for 2 days, n= 54, melphalan 140mg/m^2^. n=3 or busulphan 0.8mg/kg for 5 days, n=1) as previously reported.^1^ All patients received G-CSF-mobilized peripheral blood stem-cells from HLA A-, B-, C-, DRB1- and DQB1-matched donors and standard GvHD immunoprophylaxis with methotrexate 5mg/m^2^ on D+1, 3 and 6 and cyclosporin from Day -2 dose-titrated to keep trough levels between 200 and 300 ng/mL. Median CD34^+^ and CD3^+^ graft cell content was 5.9 (range 3.9-13.1) and 168 (range 29-458) x 10^6^/kg recipient body weight respectively. No *ex vivo* T-cell depletion of the graft or serotherapy was employed. Standard supportive care was employed including prophylactic trimethoprim/sulfamethoxazole or nebulised pentamidine to prevent *Pneumocystis jirovecii* pneumonia. Acyclovir was administered as antiviral prophylaxis. If either patient or donor was seropositive for cytomegalovirus (CMV), peripheral blood samples were monitored twice weekly for CMV re-activation by quantitative polymerase chain reaction (PCR) whilst on immunosuppression. Patients with evidence of CMV re-activation received pre-emptive ganciclovir therapy according to local protocol. T-cell lineage-specific chimerism was measured on peripheral blood samples by PCR analysis of variable number tandem repeat polymorphisms after immunomagnetic sorting of CD3^+^ T-cells. No donor lymphocyte infusions were used. Ciclosporin weaning was initiated between D+60 and D+120 according to individual chimerism status and relapse risk.

**Acute GvHD diagnosis, grading and staging and definition of relapse**

Patients were assessed for acute GvHD including both classical (pre-D+100) and late onset acute GvHD up until D+180 with organ-specific staging and overall grading according to current NIH criteria.^2^ Only Grade 2-4 acute GvHD was considered clinically significant.

**Disease Relapse**

Relapse of haematological cancer was defined as recurrence of radiological, morphological or serological disease according to standard criteria.^3-5^

**Mass cytometry panel**

The mass cytometry panel was designed and constructed using the Maxpar Panel Designer to maximize sensitivity and specificity of each antibody-lanthanide metal isotope conjugate signal on the CyTOF2 mass cytometer (Fluidigm). All antibodies were commercially sourced pre-conjugated to heavy metals except three antibodies which required custom conjugation; CCR9 which was conjugated and validated in house, and Vδ1 and Vδ2 which were supplied custom-conjugated and validated by collaborators. Antibody-metal conjugates were titrated and validated on healthy donor PBMC with metal-minus-one controls to confirm specificity of signal. Cross-validation using antibody-fluorochrome staining and conventional flow cytometry acquisition and analysis demonstrated similar frequencies of higher-order phenotypic groups and rarer populations e.g. CCR9^+^ T-cells to those using antibody-metal conjugates and mass cytometry. Panel performance was equivalent on fresh or frozen PBMC.

**Mass cytometry sample processing**

PBMC were extracted from patient and healthy donor blood by density gradient centrifugation and cryopreserved. Frozen patient samples were resuscitated, processed and run in batches of 3-5 samples on the same day. For the primary mass cytometry panel, 2.5 million viable patient PBMC and healthy donor PBMC were separately CD45-barcoded by staining with CD45-89Y and CD45-141Pr respectively, and then stained with cell surface markers (CD3, CD4, CD8a, CD45RA, CD27, CD28, CCR7, CD25, CD127, CCR5, PD1, CD19, CD16, CD56, TCRVα24, CCR9 and Vδ2). PBMC were then stained with cisplatin Cisplatin-194Pt (to facilitate exclusion of dead cells), permeabilized and stained with intranuclear markers (Tbet, FOXP3, GATA3 and RORɣT) as fixed as per manufacturer protocol. Patient PBMC were spiked with 0.5 million bar-coded healthy control PBMC and cells were tagged with a metallointercalator (Ir 191/193) to enable cellular detection, counted and suspended at 0.5 x 10^6^ cells / ml, EQ beads added at the manufacturers recommended ratio and filtered immediately prior to acquisition. A minimum of 500,000 events were acquired on the CytTOF2. For a subset of samples from D+30 the same processing and staining protocol was used with a secondary mass cytometry panel where 1μl per 1000μl of Brefeldin A solution was added for four hours prior to staining. CytofCore (0.4) in R was used to ensure channel names across mass cytometry runs were all identically named to ensure compatibility with algorithmic analysis.

**Data cleaning**

Integrated mass data (IMD) files generated by the CyTOF2 were converted to FCS 3.0 files. Arcsine transformation was used to decompress data for visualization. For data cleaning prior to high dimensional analysis (and for Boolean validation of the latter) Cytobank and Flowjo software platforms were used. Beads were gated out using Ir191 vs Ir193 plots and doublets were subsequently excluded by plotting event length vs Ir191. Live cells were captured by plotting Cisplatin 194 vs Ir191 and patient samples were resolved from healthy donor controls by plotting CD45-141Pr (spike) vs CD45-89Y (patient). CD45-postive lineage-positive live cells were used as denominator populations for both Boolean and high-dimensional analysis.

**Dimensionality reduction and data visualization**

Non-linear dimensionality reduction was used to visualize data via down-sampling as tSNE plots via the CyTOFkit platform.

**Clustering**

PhenoGraph and FlowSOM, two complimentary clustering algorithms were used to retain reproducible and stable clustering characteristics across multiple runs. PhenoGraph was utilized via a Graphical User Interface in R on the CyTOFkit platform. FlowSOM (flow or mass cytometry data and self-organising maps) was used via R in the CyTOFkit platform and FlowSOM clusters displayed on tSNE maps (CyTOFkit). The R code used for these clustering platforms in Cytofkit was as follows;

Cytofkit R

RStudio workspace code

library(“cytofkit”)

cytofkit(fcsFiles = getwd(), markers = "parameter.txt",

projectName = "cytofkit", ifCompensation = FALSE,

transformMethod = c("cytofAsinh"),

mergeMethod = c("fixed"), fixedNum = 5000,

dimReductionMethod = c("tsne"),

clusterMethods = c("RPhenoGraph", "FlowSOM"),

visualizationMethods = c("tsne"),

progressionMethod = c("NULL")

RPhenoGraph_k = 30, FlowSOM_k = 40, seed = 42,

resultDir = getwd(), saveResults = TRUE,

saveObject = TRUE, openShinyAPP = FALSE, ...)

Median marker expression of clusters was used to systematically assign clusters to cell subsets according to canonical phenotypic groups. Individual clusters were assigned to higher order phenotypic groups based on lineage markers, and subset-specific transcription factors. Single positive CD4 and CD8 T-cell clusters were grouped by naïve/memory cell differentiation (based on co-expression patterns of CD45RA and CCR7) and CD56+ NK cell clusters assigned to CD56^hi^ or CD56^dim^ groups. Correlation matrices were generated in Graphpad Prism between individual clusters. The R programme Circlize (version 0.4.5) was used to generate chordograms of correlations. Stream plots depicting the frequencies of individual immune cell populations and of higher order phenotypic groups over time were generated using DisplayR software (DisplayR, NSW, Australia).

**Boolean validation of clustering**

Frequencies of immune cell populations generated in PhenoGraph and FlowSOM clustering algorithms were validated by conventional Boolean gating analysis of mass cytometry data using Flowjo software v10.6.2 (Treestar) using canonical marker expression patterns for higher order phenotypic groups. Additionally, frequencies of immune cell populations generated in FlowSOM with significantly different frequencies in patients who subsequently developed acute GvHD and those who did not were validated by conventional Boolean gating analysis using a ‘best non-redundant phenotype’ approach.

#### Supervised learning algorithms

Two separated supervised learning algorithms, CITRUS and Diffcyt (version 1.2.10), which both employ statistical significance testing corrected for false discovery/multiple comparisons were used to validate the distribution of cell clusters generated in PhenoGraph and FlowSOM across patient groups. CITRUS combined an unsupervised clustering approach with a trained supervised model to identify significant features that predict for differences between two sample groups. Normalised FCS files were uploaded to the Cytobank server. Files from randomly selected but equal numbers of patients who did and did not subsequently develop acute GvHD were analysed as this algorithm is optimized with groups of similar size. 5000 events were down sampled from each sample for analysis, with minimum cluster size and false-discovery rate (FDR) set to 5%. All markers from the primary panel (except CD45) were used for clustering and the algorithm run five times in order to confirm and validate any findings. Diffcyt utilized FlowSOM to cluster data and edgeR to identify differential abundance between sample groups. EdgeR applied a normalisation on clusters to control for different numbers of cells in each group. Diffcyt analysis was performed on the same sample set as CITRUS. Files were loaded in to R and the Diffcyt package was set to use 5000 events / file. The output consisted of a list of the top 10 differentially abundant clusters along with adjusted p values.

**Normalization of mass cytometry data to reduce batch effects**

CD45 barcoded control PBMC from a single healthy donor were added to every patient sample to provide a consistent internal control to mitigate both variability of antibody to cell ratios in samples containing different subpopulations of cells (*intra* batch variability) and differences in staining between batches (*inter* batch variability).^6^ Control PBMC were spiked into patient samples at a ratio of 1:5 (0.5 million: 2.5 million). The two separate CD45-barcoded patient and control populations (CD45-89Y and CD45-14Pr) could be resolved easily and the addition of control cells did not impact on the frequencies of T-cell sub-populations in test samples enumerated by Boolean analysis using Flowjo software (**Supplementary Figure 1A**). A bespoke R code algorithm provided by Sophie van Gassen (now commercially available as CytoNorm) was used to normalize each patient sample to the healthy spiked controls. The ability of the algorithm to correct inter-batch variability was confirmed by assessing PhenoGraph cluster abundance in spiked healthy control cells from 30 patient samples run in 30 separate batches. Prior to normalization considerable variation in cluster abundance was seen in healthy donor cells across different samples/batches indicating a significant batch staining/acquisition effect. Normalization of spiked control cells as both source and target for the normalisation algorithm resulted in homogeneous cluster abundance distribution across batches, **Supplementary Figure 1B** without impacting on the tSNE cluster map and lineage-specific clusters, **Supplementary Figure 1C.**

**Similarity indices**

Similarity between immune cell signatures in defined patient groups using the Jaccard Similarity Index was calculated as ***(number of differentially abundant FlowSOM clusters in both groups) / (number of differentially abundant FlowSOM clusters in either group)*100.***

**Cell sorting and Suppression assays**

The following antibodies (Flurochrome, clone) were used for flow cytometric sorting of CD56^hi^ NK cells from patient samples; CD56 (PE Cy5, 5.H11, CD8 (AF700, SK1), CD45RA (BV605, HI100). DAPI was used to exclude dead cells. For suppression assays 5 x 10^4^ healthy irradiated (40Gy, Biorad RS2000 X-ray irradiator) allogeneic donor PBMC stimulators were co-cultured for 8 days with equal numbers of CFSE-labelled patient PBMC responders in 96 well round-bottomed plates with or without addition of 5 x 10^4^ patient flow-sorted CD56^hi^ cells. Technical replicates and autologous controls were used. On day 8, the cells were stained using cell surface and intracellular markers including the following antibodies (Flurochrome, clone); CD4 (PerCP-Cy5.5, RPA-T4), CD8 (AF700, SK1), CD3 (BV786, SK7), CD56 (PE Cy5, 5.1H11) Perforin (BV421, B-D48). All antibodies were from Biolegend. Zombie red was used as viability marker in the suppression assays. Brefeldin was added 3 hours prior to staining and cells were acquired on an LSRFortessa (Becton Dickenson). Data was analysed using FlowJo version 10 (Treestar). Alloproliferation was calculated as (% CFSE^lo^ in allostimulated cells – % CFSE^lo^ in autologous control). Allosuppression (%) was calculated as 100 x (1-% alloproliferation of patient sample with putified CD56hi cells /% alloproliferation of patient sample).

**Tumour-associated antigen (TAA)-specific T-cell enumeration**

TAA-specific T-cell staining using HLA-A2-restricetd dextramers was optimized using healthy donor PBMC sourced from NHSBT, HLA A2-negative PBMC were used as negative controls. HLA-A201^+^ patient samples and healthy control PBMC were resuscitated as previously described. Cell counts and viability were assessed and only samples that had >80% live cells were used. 10µl WT1-PE and PR1-APC were added to two million live cells from each sample and incubated in the dark for 10 minutes at room temperature. Immediately subsequent to Dextramer staining anti-CD3 (clone SK7, BV785, Biolegend) and anti-CD8 (clone SK1, FITC, Biolegend) were added and samples were incubated on ice for 20 minutes in the dark. Cells were washed and DAPI added prior to acquisition on a BD LSRFortessa cell analyser and analysed using FlowJo version 10 (Treestar) software. A minimum of 50 CD3^+^ CD8^+^ Dextramer^+^ events was acquired wherever possible to maintain an assay co-efficient of variance of < 15%. For each sample the true percentage positive events were calculated by subtracting the number of false positive events obtained in the HLA-A2 negative healthy control.

**Measurement of serum protein biomarkers**

Patient serum samples were centrifuged prior to use, diluted using Calibrator Diluent RD6-52 and 50µL of standard or sample was added in to each well. The diluted microparticle cocktail was added to each well of a microplate, covered with foil and incubated for 2 hours on a horizontal microplate shaker at 800rpm then washed three times. A magnet was applied to the bottom of the microplate well and the liquid was removed, before each well was refilled with wash buffer and 50 μL of diluted Biotin-Antibody Cocktail was added to each well, covered and incubated for 1 hour at room temperature on the shaker. The wash step was repeated three times. 50 μL of diluted Streptavidin-PE was added to each well, covered and incubated for 30 minutes at room temperature on the shaker set at 800rpm. The wash step was then repeated as described three times. The microparticles were re-suspended, the plate was incubated for 2 minutes on the shaker set at 800rpm. The plate was incubated overnight at 4 degrees and then read the next day using the Luminex analyser. (^*p*),^7^ a surrogate predictor for aGvHD was calculated as log10[–log10(1 – ^*p*)] = –11.263 + 1.844(log10ST2) + 0.577(log10REG3α).

**References**

1.	Davies JK, Taussig D, Oakervee H, et al. Long-term survival with low toxicity after allogeneic transplantation for acute myeloid leukaemia and myelodysplasia using non-myeloablative conditioning without T cell depletion. *British journal of haematology.* 2013;162(4):525-529.

2.	Filipovich AH, Weisdorf D, Pavletic S, et al. National Institutes of Health consensus development project on criteria for clinical trials in chronic graft-versus-host disease: I. Diagnosis and staging working group report. *Biology of blood and marrow transplantation : journal of the American Society for Blood and Marrow Transplantation.* 2005;11(12):945-956.

3.	Cheson BD, Horning SJ, Coiffier B, et al. Report of an international workshop to standardize response criteria for non-Hodgkin's lymphomas. NCI Sponsored International Working Group. *J Clin Oncol.* 1999;17(4):1244.

4.	Durie BG, Harousseau JL, Miguel JS, et al. International uniform response criteria for multiple myeloma. *Leukemia.* 2006;20(9):1467-1473.

5.	Dohner H, Estey EH, Amadori S, et al. Diagnosis and management of acute myeloid leukemia in adults: recommendations from an international expert panel, on behalf of the European LeukemiaNet. *Blood.* 2010;115(3):453-474.

6.	Kleinsteuber K, Corleis B, Rashidi N, et al. Standardization and quality control for high-dimensional mass cytometry studies of human samples. *Cytometry A.* 2016;89(10):903-913.

7.	Hartwell MJ, Ozbek U, Holler E, et al. An early-biomarker algorithm predicts lethal graft-versus-host disease and survival. *JCI Insight.* 2017;2(3):e89798.

8.	Sallusto F, Lenig D, Forster R, Lipp M, Lanzavecchia A. Two subsets of memory T lymphocytes with distinct homing potentials and effector functions. *Nature.* 1999;401(6754):708-712.

9.	Koch S, Larbi A, Derhovanessian E, Ozcelik D, Naumova E, Pawelec G. Multiparameter flow cytometric analysis of CD4 and CD8 T cell subsets in young and old people. *Immun Ageing.* 2008;5:6.

**Table S1 Primary mass cytometry panel for all samples**

| Class | Marker | Class | Marker |
| --- | --- | --- | --- |
| Lineage |  | **Naïve/Memory** | CD45RA |
| Haematopoietic | CD45 |  | CCR7 |
| T-cell | CD3 |  | CD27 |
| T-cell subset | CD4 |  | CD28 |
|  | CD8 |  |  |
| γδ T-cell | Vδ1 | **Chemotaxis** | CCR5 |
|  | Vδ2 |  | CCR7 |
| B cell | CD19 |  | CCR9 |
| NK cell | CD56 |  |  |
|  | CD16 | **Function** |  |
| iNKT | TCRVα24 | Treg | FOXP3 |
|  |  | Th1/Tc1 | T-bet |
| Viability | Cisplatin | Th2/Tc2 | GATA 3 |
|  |  | Th17 | RORγT |
| Activation | CD25 | **Exhaustion** | PD1 |
|  | CD127 |  |  |

**Table S2 Antibody-metal conjugates for primary mass cytometry panel**

| Marker | CyTOF2 Metal | Antibody Clone |
| --- | --- | --- |
| CD3 | 154Sm | UCHT1 |
| CD4 | 145Nm | RPAT4 |
| CD8α | 146Nd | RPAT8 |
| CD45RA | 153Eu | HI100 |
| CD27 | 155Gd | L128 |
| CD28 | 160GdM | CD28.2 |
| CCR7 | 159Tb | G043H7 |
| Tbet | 161Dy | 4B10 |
| GATA3 | 167Er | TWAJ |
| RORγT | 168Er | NR1F3 |
| CD25 | 149Sm | 2A3 |
| CD127 | 165Ho | A019D5 |
| FOXP3 | 162Dy | PCH101 |
| CCR5 | 156Gd | NP=6G4 |
| CCR9 | 169Tm | Lo53E8 |
| PD-1 | 175Lu | EH12.2H7 |
| CD19 | 142Nd | H1B19 |
| CD16 | 209Bi | 3G8 |
| CD56 | 176Yb | NCAM16.2 |
| TCRvα24Ja18 | 170Eu | 6B11 |
| Vδ1 | 173Yb | TS8.2 |
| Vδ2 | 143Nd | B6 |
| CD45 | 89-Y and 141Pr | HI30 |
| Cisplatin | 194Pt | N/A |

**Table S3 Secondary mass cytometry panel for subset of samples**

| Class | Marker | Class | Marker |
| --- | --- | --- | --- |
| Lineage |  | **Naïve/Memory** | CD45RA |
| Haematopoietic | CD45 |  | CCR7 |
| T-cell | CD3 |  | CD27 |
| T-cell subset | CD4 |  | CD28 |
| γδ T-cell | Vδ1 | **Chemotaxis** | CCR5 |
|  | Vδ2 |  | CCR7 |
| B cell | CD19 |  |  |
| NK cell | CD56 | **NK receptors** | NKG2A |
|  | CD16 |  | NKp46 |
|  |  | **Function** |  |
| iNKT | TCRVα24 | Th1/Tc1/Cytolytic | Tbet |
|  |  | Cytolytic | Perforin |
|  |  | Cytolytic | IFN-γ |
|  |  | Suppression | TGF-β |
|  |  | Suppression | IL-10 |
| Viability | Cisplatin |  |  |

**Table S4 Antibody-metal conjugates for secondary mass cytometry panel**

| Marker | CyTOF2 Metal | Antibody Clone |
| --- | --- | --- |
| CD3 | 154Sm | UCHT1 |
| CD4 | 145Nm | RPAT4 |
| CD45RA | 153Eu | HI100 |
| CD27 | 155Gd | L128 |
| CD28 | 160GdM | CD28.2 |
| CCR7 | 159Tb | G043H7 |
| Tbet | 161Dy | 4B10 |
| CCR5 | 156Gd | NP=6G4 |
| CD19 | 142Nd | H1B19 |
| CD16 | 209Bi | 3G8 |
| CD56 | 176Yb | NCAM16.2 |
| TCRvα24Ja18 | 170Eu | 6B11 |
| Vδ1 | 173Yb | TS8.2 |
| Vδ2 | 143Nd | B6 |
| CD45 | 89-Y and 141Pr | HI30 |
| Cisplatin | 194Pt | N/A |
| NKG2A | 169Tm | Z199 |
| NKp46 | 162Dy | BAB281 |
| Perforin | 175Lu | B-D48 |
| TGF-β | 163Dy | TW46H10 |
| IFN-γ | 165Ho | B27 |
| Il-10 | 166Fr | JES3-9D7 |

**Table S5 Demographics, donor details and transplant conditioning for AHST patients**

| UPN | Gender | Age  (yrs) | Diag | Donor | Donor gender | Sex m/m* | Status | CMV risk | ABO m/m | Conditioning |
| --- | --- | --- | --- | --- | --- | --- | --- | --- | --- | --- |
| 1 | M | 60 | MDS | MUD | F | Yes | PR>1 | High | Minor | Flu-Cy |
| 2 | F | 54 | NHL | MUD | F | No | CR>1 | High | None | Flu-Cy |
| 3 | M | 56 | NHL | MSD | F | Yes | CR1 | Low | None | Flu-Cy |
| 4 | F | 49 | AML | MUD | M | No | CR1 | High | Major | Flu-Cy |
| 5 | M | 74 | MDS | MUD | F | Yes | MDS | High | None | Flu-Cy |
| 6 | F | 32 | AML | MUD | M | No | CR1 | High | Minor | Flu-Cy |
| 7 | M | 55 | MF | MUD | M | No | MF | Low | minor | Flu-Mel |
| 8 | M | 53 | AML | MUD | M | No | CR>1 | High | Major | Flu-Cy |
| 9 | M | 40 | AML | MSD | M | No | CR>1 | High | None | Flu-Cy |
| 10 | F | 63 | AML | MUD | M | No | CR1 | High | None | Flu-Cy |
| 11 | M | 59 | AML | MUD | M | No | CR>1 | High | Major | Flu-Cy |
| 12 | M | 51 | ALL | MUD | M | No | CR>1 | High | None | Flu-Cy |
| 13 | F | 46 | MM | MSD | F | No | PR1 | Low | Minor | Flu-Cy |
| 14 | M | 67 | ALL | MSD | F | Yes | CR1 | Low | None | Flu-Cy |
| 15 | M | 37 | AML | MUD | M | No | CR1 | High | None | Flu-Cy |
| 16 | F | 65 | AML | MUD | M | No | CR1 | High | None | Flu-Cy |
| 17 | M | 55 | ALL | MSD | M | No | CR1 | High | None | Flu-Cy |
| 18 | M | 51 | MM | MSD | F | Yes | PR1 | High | None | Flu-Cy |
| 19 | F | 31 | AML | MUD | M | No | PR>1 | High | None | Flu-Cy |
| 20 | F | 60 | AML | MUD | M | No | CR1 | Low | Minor | Flu-Cy |
| 21 | F | 67 | AML | MUD | M | No | CR1 | Low | None | Flu-Cy |
| 22 | F | 67 | AML | MUD | M | No | CR1 | High | Major | Flu-Cy |
| 23 | M | 49 | MDS | MSD | F | Yes | MDS | High | Minor | Flu-Cy |
| 24 | M | 26 | NHL | MSD | F | Yes | CR1 | Low | Major | Flu-Cy |
| 25 | M | 54 | AML | MUD | M | No | CR1 | High | Major | Flu-Mel |
| 26 | F | 38 | MM | MUD | M | No | PR>1 | Low | None | Flu-Cy |
| 27 | F | 59 | AML | MUD | M | No | CR1 | Low | Minor | Flu-Cy |
| 28 | F | 66 | NHL | MUD | F | No | CR1 | Low | None | Flu-Cy |
| 29 | M | 64 | AML | MSD | M | No | CR>1 | Low | None | Flu-Cy |
| 30 | M | 41 | AML | MUD | M | No | CR1 | High | None | Flu-Cy |
| 31 | M | 65 | TPLL | MUD | M | No | CR1 | Low | minor | Flu-Bu |
| 32 | M | 49 | AML | MUD | M | No | CR>1 | High | None | Flu-Cy |
| 33 | M | 70 | AML | MUD | M | No | CR1 | High | Major | Flu-Cy |
| 34 | M | 68 | AML | MUD | M | No | CR1 | High | Minor | Flu-Cy |
| 35 | M | 77 | AML | MSD | M | No | CR>1 | High | None | Flu-Cy |
| 36 | F | 33 | AML | MUD | M | No | CR1 | High | Minor | Flu-Cy |
| 37 | M | 54 | MM | MSD | M | No | PR | Low | None | Flu-Cy |
| 38 | M | 65 | MDS | MUD | M | No | PR1 | High | None | Flu-Cy |
| 39 | M | 37 | AML | MUD | M | No | CR1 | High | None | Flu-Cy |
| 40 | F | 36 | AML | MSD | M | No | CR1 | High | None | Flu-Cy |
| 41 | M | 54 | NHL | MUD | M | No | CR1 | High | Major | Flu-Cy |
| 42 | M | 73 | CMML | MUD | M | No | PR>1 | High | Major | Flu-Cy |
| 43 | M | 32 | HL | MUD | M | No | PR1 | Low | None | Flu-Cy |
| 44 | M | 46 | MDS | MSD | M | No | MDS | High | Major | Flu-Cy |
| 45 | F | 66 | AML | MUD | M | No | CR1 | High | Minor | Flu-Cy |
| 46 | M | 73 | MDS | MSD | M | No | MDS | High | Major | Flu-Cy |
| 47 | M | 53 | MM | MSD | F | Yes | CR1 | Low | None | Flu-Cy |
| 48 | F | 51 | AML | MUD | M | No | CR1 | Low | None | Flu-Cy |
| 49 | F | 46 | NHL | MSD | F | No | PR>1 | High | None | Flu-Cy |
| 50 | F | 60 | AML | MUD | M | No | CR1 | High | None | Flu-Cy |
| 51 | M | 60 | AML | MSD | M | No | CR1 | High | None | Flu-Cy |
| 52 | M | 74 | AML | MUD | M | No | CR1 | Low | None | Flu-Cy |
| 53 | M | 67 | CMML | MUD | M | No | CMML1 | High | None | Flu-Cy |
| 54 | F | 53 | MM | MSD | M | No | PR1 | High | None | Flu-Cy |
| 55 | F | 26 | ALL | MUD | M | No | CR1 | Low | Major | Flu-Mel |
| 56 | M | 55 | CMML | MSD | M | No | CR1 | Low | None | Flu-Cy |
| 57 | F | 26 | ALL | MUD | M | No | CR1 | Low | Minot | Flu-Cy |
| 58 | M | 48 | CMML | MSD | F | Yes | CMML1 | High | None | Flu-Cy |

UPN, Unique patient number; M, male, F, female, yrs, years, AML, acute myeloid leukemia; MDS, myelodysplasia; ALL, Acute lymphoblastic leukemia; CMML, chronic myelomonocytic leukemia; NHL, non-hodgkin lymphoma; MCL, Mantle cell lymphoma; HL, Hodgkin lymphoma; MF, myelofibrosis; MM, multiple myeloma; MUD, matched (HLA-A, B, C, DR and DQ) unrelated donor; MSD, matched (HLA-A, B, C, DR) sibling donor; PR, partial remission; CR, complete remission; Low denotes CMV seronegative donor and recipient, High denotes any other combination; Flu, Fludarabine; Cy, cyclophosphamide; Mel, Melphalan; Bu, Busulphan

* sex mismatch defined as female donor and male recipient

**Table S6 Pre-transplant factors and frequencies of D+30 Flowsom CD56^hi^ 6 and CD4 EM2 clusters**

| Variable | Group | FlowSOM CD56^hi^ 6  median frequency (range) | P value (MWT) | FlowSOM CD4 EM2  Median frequency (range) | P value (MWT) |
| --- | --- | --- | --- | --- | --- |
| Recipient Age | Below median | 2.2 (0.04-14.54) | 0.61 | 1.3 (0.16-7.3) | 0.21 |
|  | Above Median | 2.92 (0.02-13.92) |  | 0.92 (0.18-3.02) |  |
| Donor | Family | 3.1 (0.66-9.62) | ***0.02*** | 1.49 (0.42-6.02) | 0.18 |
|  | Unrelated | 1.59 (0.02-14.54) |  | 1.15 (0.16-7.38) |  |
| CD34 count | Below median | 1.09 (0.04-3.22) | 0.46 | 1.02 (0.16-1.5) | 0.39 |
|  | Above median | 1.39 (0.26-4.08) |  | 2.36 (0.30-7.38) |  |
| CD3 count | Below median | 2.45 (0.04-3.90) | 0.28 | 0.45 (0.16-1.50) | 0.07 |
|  | Above median | 0.48 (0.16-4.08 |  | 1.12 (0.54-7.38) |  |

**Figure S1**

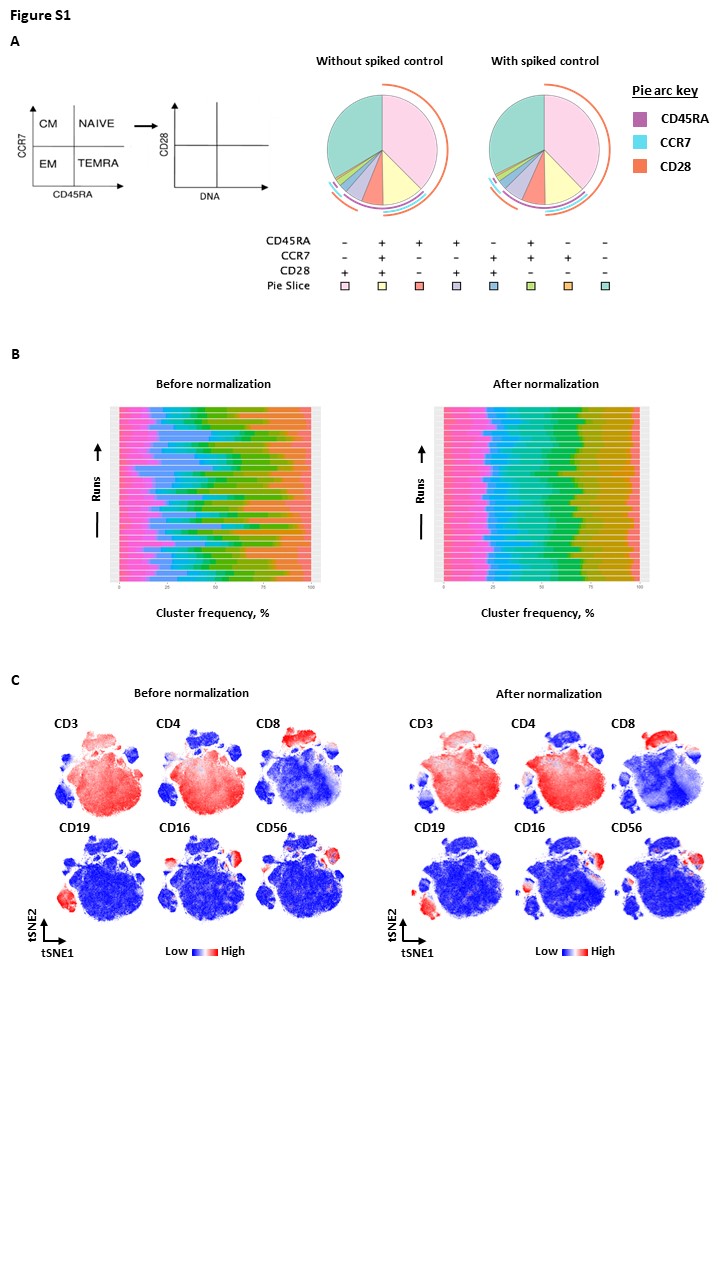

**Figure S1 Quality control of samples with barcoded healthy spiked controls and normalization algorithm**

**A** Proportions of naïve and effector memory CD8 T-cells defined by co-expression patterns of CD45RA and CCR7^8^ and CD28^9^ in healthy donor PBMCs assessed by mass cytometry and Boolean analysis were not affected by the addition of CD45-barcoded control PBMC from a different healthy donor.

**B** Batch effects were significant but effectively reduced by normalization. Frequencies of lineage-specific immune cell clusters generated by PhenoGraph in spiked control PBMC from the same donor run in 30 separate batch assays are shown before and after normalization.

**C** High dimensional visualization of immune cell populations in healthy donor PBMCs assessed by mass cytometry remained intact after normalization to CD45-barcoded control PBMC from a different healthy donor. tSNE plots with expression of lineage markers are shown before and after normalization.

**Figure S2**

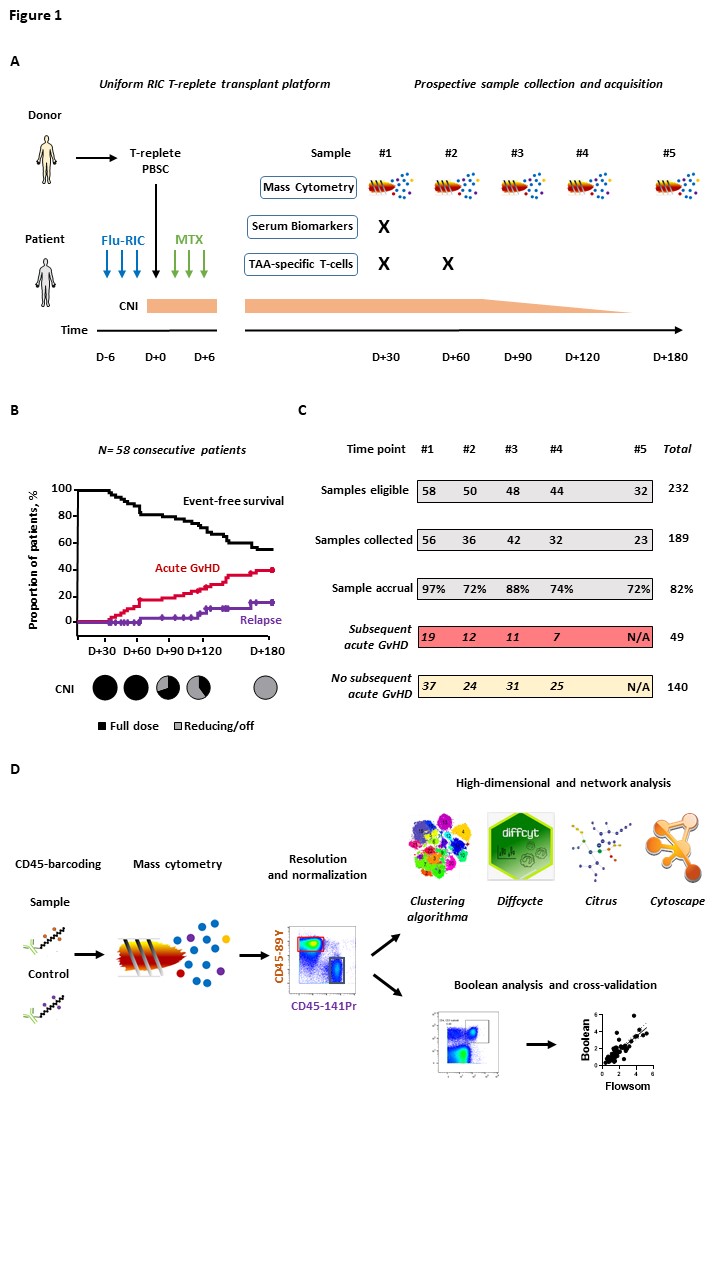

**Figure S2 Study design, sample accrual and analysis pipeline**

**A** Transplant platform, post-transplant immunoprophylaxis and sampling time-points. RIC, reduced-intensity conditioning; PBSC, peripheral blood stem-cells; Flu, Fludarabine; MTX, Methotrexate; CNI, calcineurin inhibitor; TAA, Tumour-associated antigen

**B** Proportion of patients eligible for immune reconstitution sample analysis over time after AHST. Figure depicts event-free survival (no acute GvHD, relapse or treatment-related mortality) and CI of aGvHD and relapse up to D+180. Patients were eligible for immune reconstitution sample analysis only if they remained event-free by each sampling time-point.

**C** Numbers of patients eligible for immune reconstitution sample analysis, sample accrual and numbers of samples from patients who subsequently developed or remained free of aGvHD to D+180 at each time-point.

**D** Pipeline for sample acquisition for single cell mass cytometry with CD45 barcoded internal controls and multi-algorithmic analysis.

**Figure S3**

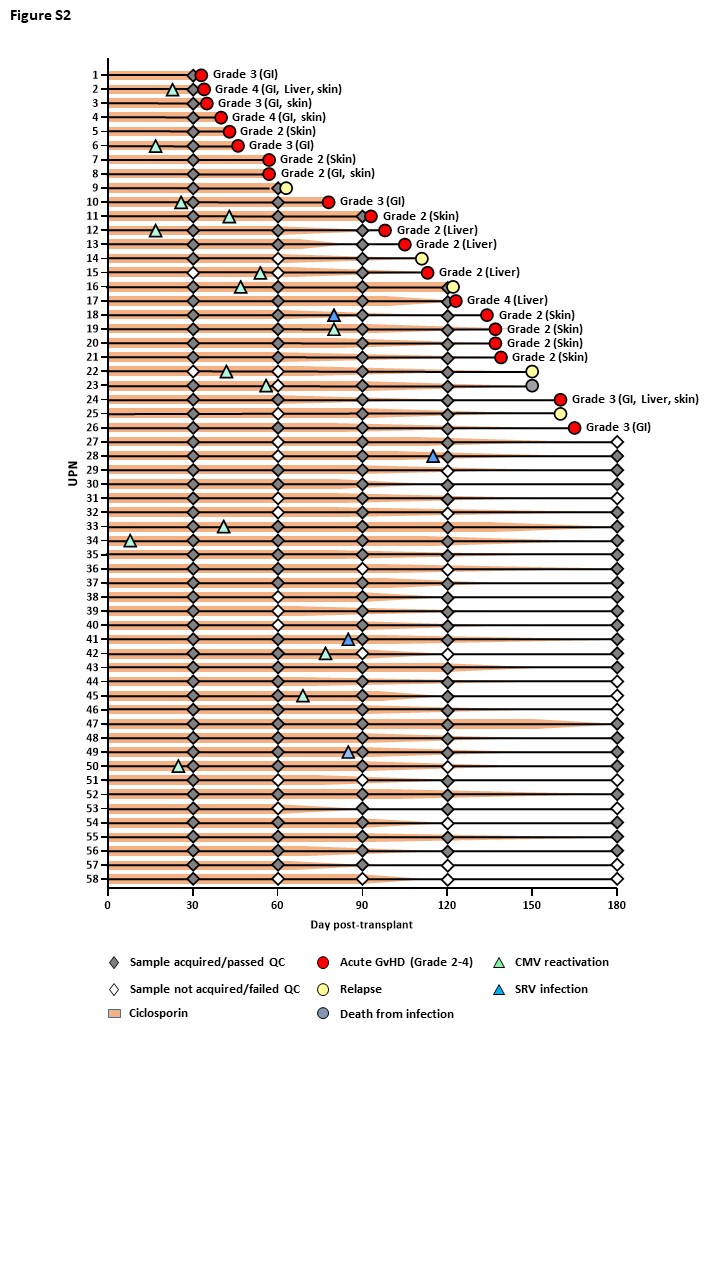

**Figure S3 Swim plot denoting samples analysed and clinical outcomes of individual patients in cohort.**

Timing of withdrawal of calcineurin inhibition, overall grade and organ involvement of acute GvHD and episodes of clinically significant CMV reactivation and seasonal respiratory virus (SRV) infection are also depicted (at the time of start of reactivation/infection). GI, gastro-intestinal.

**Figure S4**

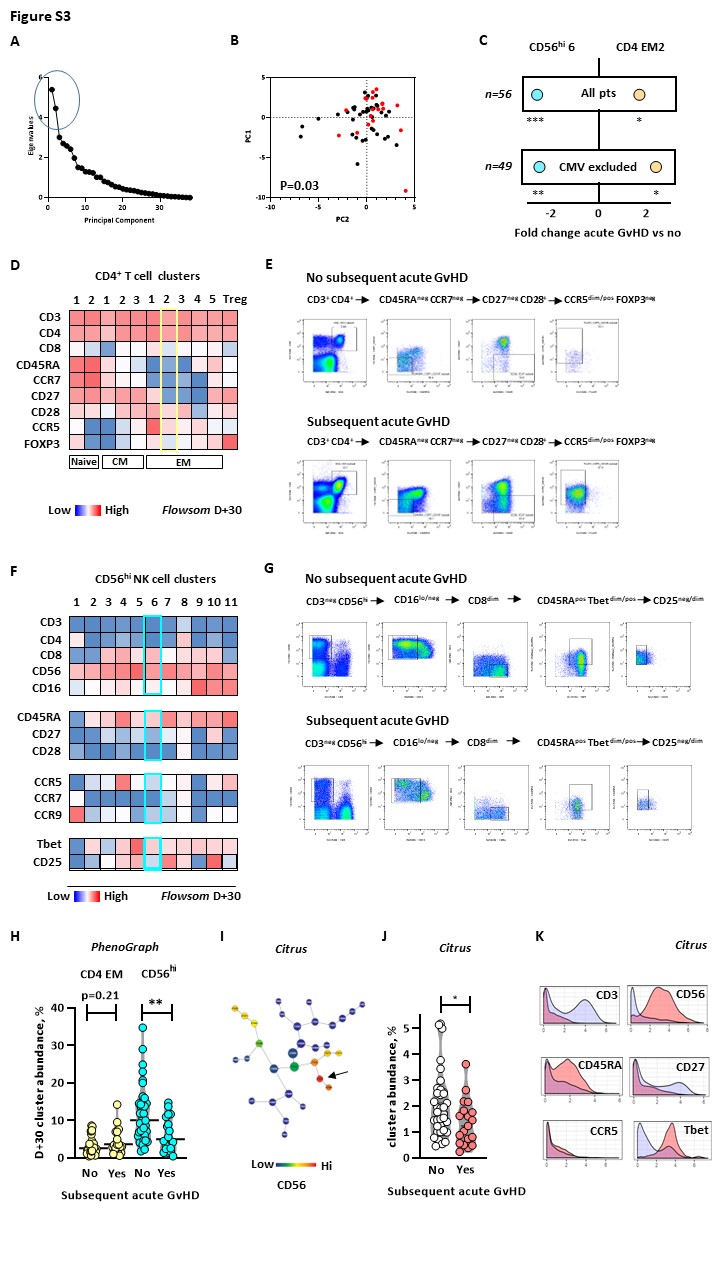

**Figure S4 Principal components analysis of FlowSOM clusters at D+30 after AHST and orthogonal validation of differential cell clusters predictive of subsequent acute GvHD**

**A** Scree plot depicted the two principal components (PC1 and PC2) with the largest Eigenvalues (circled)

**B** Bivariate plot showing PC1 and PC2 values for all patient samples with samples from patients who subsequently developed acute GvHD shown in red. P value is for Fishers exact test for distribution of samples in the PC1^hi^ PC2^hi^ quadrant compared to samples in the remaining 3 quadrants.

**C** Figure depicts fold change in frequency of CD4 EM2 T-cell and CD56^h^i 6 NK cell FlowSOM clusters in patients who did and did not subsequently develop acute GvHD in the whole patient cohort (upper) and after exclusion on patients who had reactivated and cleared CMV prior to D+30 (lower). P values are for MWT. * <0.05, ** p<0.01, *** p< 0.001

**D** Heat map depicting median marker expression for markers used to identify the closest non-redundant phenotype to the differential CD4 EM2 cluster identified using the FlowSOM algorithm (yellow box). The closest non-redundant phenotype was CD3^+^ CD4^+^ CD45RA^neg^ CCR7^neg^ CD27^neg^ CD28^pos^ CCR5^dim/pos^ FOXP3^neg^

**E** Boolean gating approach to enumerate live cells with the closest non-redundant phenotype to FlowSOM CD4 EM2 cluster in representative patients who did not (upper) and did (lower) subsequently develop acute GvHD.

**F** Heat map depicting median marker expression for markers used to identify the closest non-redundant phenotype to the differential CD56^hi^ 6 cluster identified using the FlowSOM algorithm (light blue box). The closest non-redundant phenotype was CD3^neg^ CD56^hi^ CD16^dim/neg^ CD4^neg^ CD8^dim^ CD45RA^+^ CCR5^neg^ Tbet^dim/pos^ CD25^dim/neg^

**G** Boolean gating approach to enumerate live cells with the closest non-redundant phenotype to FlowSOM CD56^hi^ 6 cluster in representative patients who did not (upper) and did (lower) subsequently develop acute GvHD.

**H** Differential CD4 EM T-cell and CD56^hi^ NK cell clusters were also identified in patients who subsequently developed acute GvHD using PhenoGraph clustering, with the same vector and similar magnitude of change to FlowSOM clustering.

**I-K** A differential CD56^hi^ cell cluster were also identified in patients who subsequently developed acute GvHD using alternative high dimensional analysis algorithm CITRUS. A CTRUS tree plot is shown with a differential CD56^hi^ cluster with significantly higher frequency in patients who did not develop acute GvHD identified in CITRUS analysis, arrowed (**B**). The frequency (**C**) and phenotype (**D**) of this differential CD56^hi^ CITRUS cluster was very similar to the CD56^hi^ clusters identified by the Flowsom and PhenoGraph models.

**Figure S5**

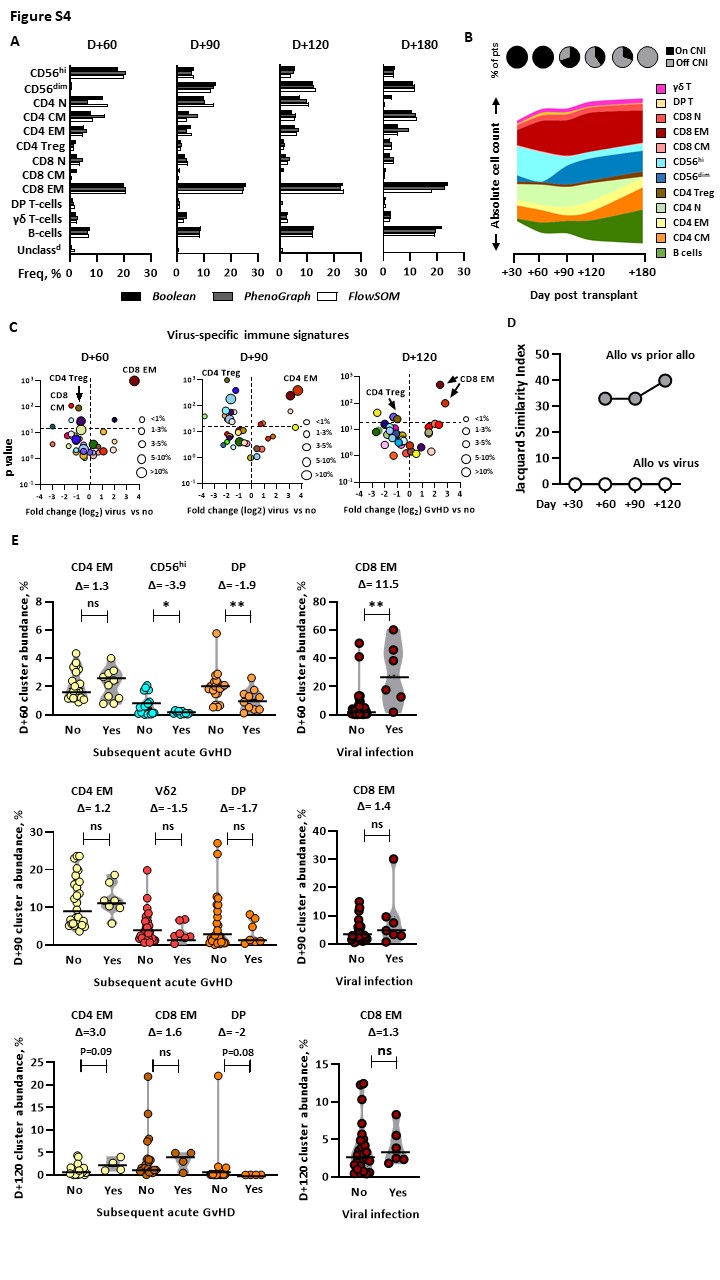

**Figure S5 The immune cell landscape at later time points after AHST and orthogonal validation of immune cell clusters predictive of subsequent acute GvHD**

**A** Frequencies of NK, T- and B-cell higher order phenotypic groups (HOPG) remained closely concordant using either PhenoGraph or FlowSOM clustering algorithms or Boolean gating analysis at later time-points after AHST.

**B** The CD56^hi^ NK cell compartment contracted whereas the CD56^dim^ NK, CD4 and CD8 EM T-cell and B-cell compartments expanded over time concurrent with the withdrawal of calcineurin inhibitor therapy. Figure depicts a stream plot of the absolute count (cells/μL) of summated FlowSOM clusters within each HOPG at each time-point. Pie charts depict the proportion of patients on or off/weaning calcineurin inhibition

**C** The virus-associated immune profile characterized by selective expansion of CD8 EM T-cell clusters in patients was maintained over time after AHST. Volcano plots depict log_2_ fold change in individual cluster abundance in patients with/without clinically significant viral reactivation/infection. Clusters are derived from FlowSOM analysis. To reduce any confounding effect of alloreactive immune signatures patients who developed aGvHD within 30 days of each time-point are excluded.

**D** Allospecific immune signatures evolved over time and remained distinct from viral immune signatures. Jaccard Index Similarity Scores are shown for allospecific signatures and allospecific signatures for the prior time-point (grey) and allospecific and viral signatures at the same time-points (white).

**E** Validation of differential FlowSOM clusters in immune signatures at later time-points after AHST using PhenoGraph analysis. Left hand plots show the frequencies of PhenoGraph clusters with analogous phenotype to the differential FlowSOM clusters in patients who subsequently developed acute GVHD. Patients with clinically significant viral reactivation/infection were excluded. Right hand plots show frequencies of PhenoGraph cluster with analogous phenotype to the largest differential FlowSOM CD8 EM T-cell cluster in patients with viral reactivation/infection. Horizontal lines are medians, P values are for MWT, Δ, fold change, ns, not significant, * p<0.05, ** p<0.001

**Figure S6**

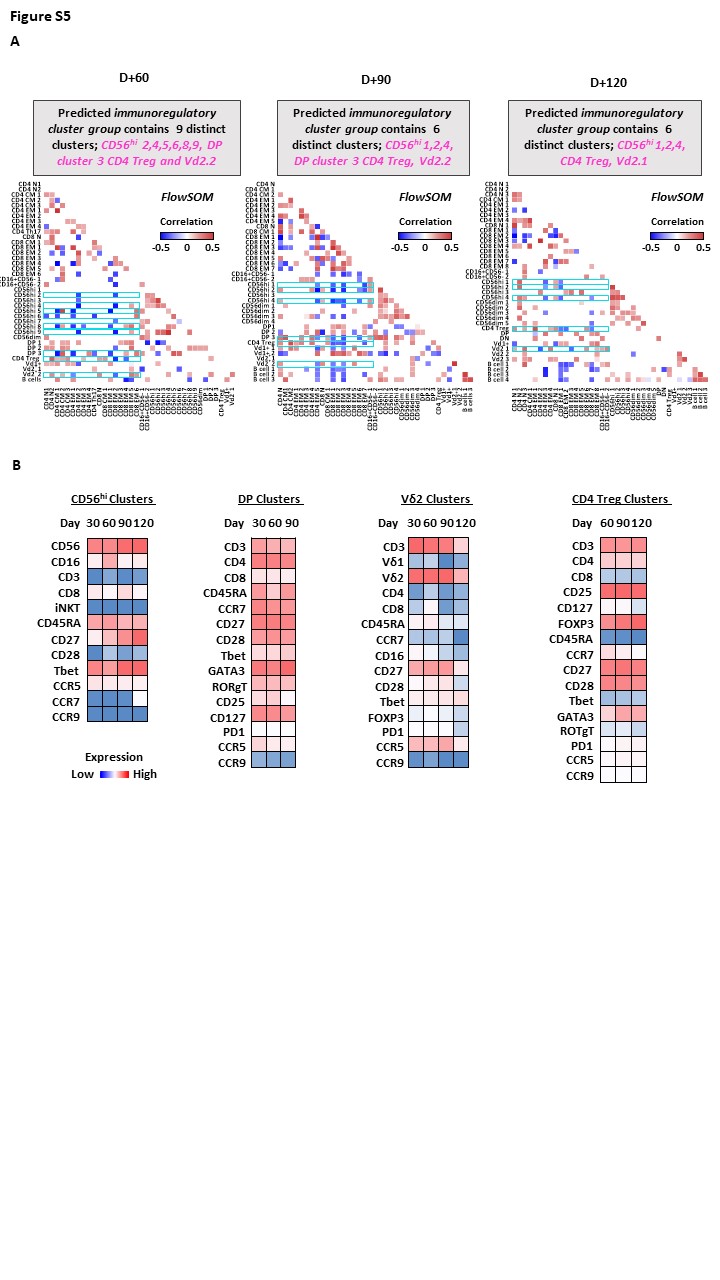

**Figure S6 Cellular populations within the predicted IRCG at later time points after AHST**

**A** Correlation analysis identified clusters with known immunoregulatory function which were significantly negatively correlated with 2 or more CD4 and/or CD8 EM T-cell clusters at later time-points after AHST. These clusters, highlighted with blue boxes, formed the predicted immunoregulatory cluster group. Correlation matrices with heat maps depict only significant (p<0.05) Spearman correlations with r > modulus 0.30. Clusters were generated using Flowsom.

**B** Heat map depicting phenotype of major IRCG component clusters over time after AHST. Median expression levels for cells in each cluster are shown. Clusters were generated using Flowsom.
